# Improving functional recovery after severe spinal cord injury by a noninvasive dual functional approach of neuroprotection and neuromodulation

**DOI:** 10.1101/2022.02.14.478109

**Authors:** Yanming Zuo, Jingjia Ye, Wanxiong Cai, Binjie Guo, Xiangfeng Chen, Lingmin Lin, Shuang Jin, Hanyu Zheng, Ao Fang, Xingran Qian, Zeinab Abdelrahman, Zhiping Wang, Zhipeng Zhang, Bin Yu, Xiaosong Gu, Xuhua Wang

## Abstract

Despite tremendous unmet medical needs, there is no effective pharmacological treatment to promote functional recovery after spinal cord injury (SCI). Although multiple pathological events have been implicated in SCI, the development of a noninvasive pharmacological approach to simultaneously target the different mechanisms involved in SCI remains a formidable challenge. In this study, we report the development of a noninvasive nanodrug delivery system that consists of ROS-responsive amphiphilic copolymers and an encapsulated neurotransmitter-conjugated KCC2 agonist. We show that upon intravenous administration, the nanodrugs were able to enter the injured spinal cord due to blood spinal cord barrier disruption and ROS-responsive disassembly. Remarkably, once in the injured spinal cord, these nanodrugs exhibited dual functions: scavenging ROS accumulated in the lesion to protect spared connections and increasing neuronal excitability in the injured spinal cord through targeted delivery of the KCC2 agonist to inhibitory neurons. Thus, the noninvasive treatment led to significant functional recovery in the rats with contusive SCI. Together, these findings provide a much-needed translational pharmacological approach for treating severe SCI.

## Introduction

It is known that most of human patients of spinal cord injury (SCI) are anatomically incomplete, meaning that some connections in neuronal circuits above and below the lesion are spared. However, at least two major processes prevent these spared connections from functioning. First, cell death and blood vessel disruption resulting from traumatic injury trigger inflammation and the production of cytotoxic factors such as reactive oxygen species (ROS) through a process termed secondary injury, further damaging the spinal connections that were spared after the primary injury and exacerbating functional deficits^1, 2^. Intensive efforts have been made in the past decade to develop protective treatment strategies that can rescue spared spinal connections, such as the transplantation of neuroprotective materials^3–5^. However, these invasive procedures might further damage the injured spinal cord and result in unpredictable side effects. Second, SCI triggers massive alterations in excitability, disrupting the overall balance of neural circuits in the injured spinal cord^6–8^. One demonstrated mechanism that underlies these changes is injury-induced downregulation of the expression of KCC2, a neuron-specific potassium chloride cotransporter, in neurons in injured spinal cord^6, 9^. Previous studies have reported that inhibition of inhibitory, but not excitatory, interneurons via genetic activation of KCC2 can increase overall excitability and transform initially dormant relay or endogenous spinal circuits into a functional state after SCI^6, 9^. However, it is yet not clear whether this strategy is efficacious in clinically translatable SCI models. In addition, the development of a noninvasive pharmacological approach to selectively target inhibitory neurons and alter their activity remains the bottleneck for the clinical application of this strategy. Thus, an ideal therapeutic strategy should target these mechanisms simultaneously in a noninvasive manner.

Noninvasive methods, such as the use of polymeric nanoparticles encapsulating drugs, have been shown to be highly efficient because unlike conventional small-molecule drugs, these agents are not affected by poor stability, short blood retention, and mistargeting problems^10, 11^. However, a hurdle to the application of these treatments^12–15^ for CNS diseases such as SCI is that these agents cannot easily enter the CNS because of blood brain barrier (BBB) or brain spinal cord barrier (BSCB). In this regard, a recent study showed that BSCB disruption permits intravenously injected AAV9 to enter the injured spinal cord9. Although the underlying mechanism of injury-induced BSCB alterations remains unknown, this finding suggests the exciting possibility of designing relevant agents that can enter the spinal cord lesion.

Another remaining issue is selective targeting of relevant neuronal types. Disguising drugs as neurotransmitters can promote their delivery into neurons that synthesize these neurotransmitters^16–18^. Thus, we hypothesized that a similar strategy could be used to deliver drugs that target inhibitory interneurons. Specifically, we conjugated the drug CLP-257, a verified KCC2 activator that reduces the intracellular chloride concentration and decreases neuronal excitability^19–21^, to γ-aminobutyric acid (GABA), which has been demonstrated to play important roles in motor control^16, 22–24^, or dopamine^25–27^ (Fig. 1a). To facilitate noninvasive administration and ROS-responsive release, an amphiphilic block copolymer with a PEG-based hydrophilic segment and a hydrophobic segment with boron-based ROS scavengers was designed and synthesized to encapsulate the hydrophobic prodrugs^28–31^ (Fig. 1a). Such micelle-based nanodrugs could be easily intravenously injected, and ROS responsively release prodrugs at injury sites^30, 32, 33^.

**Fig. 1.**
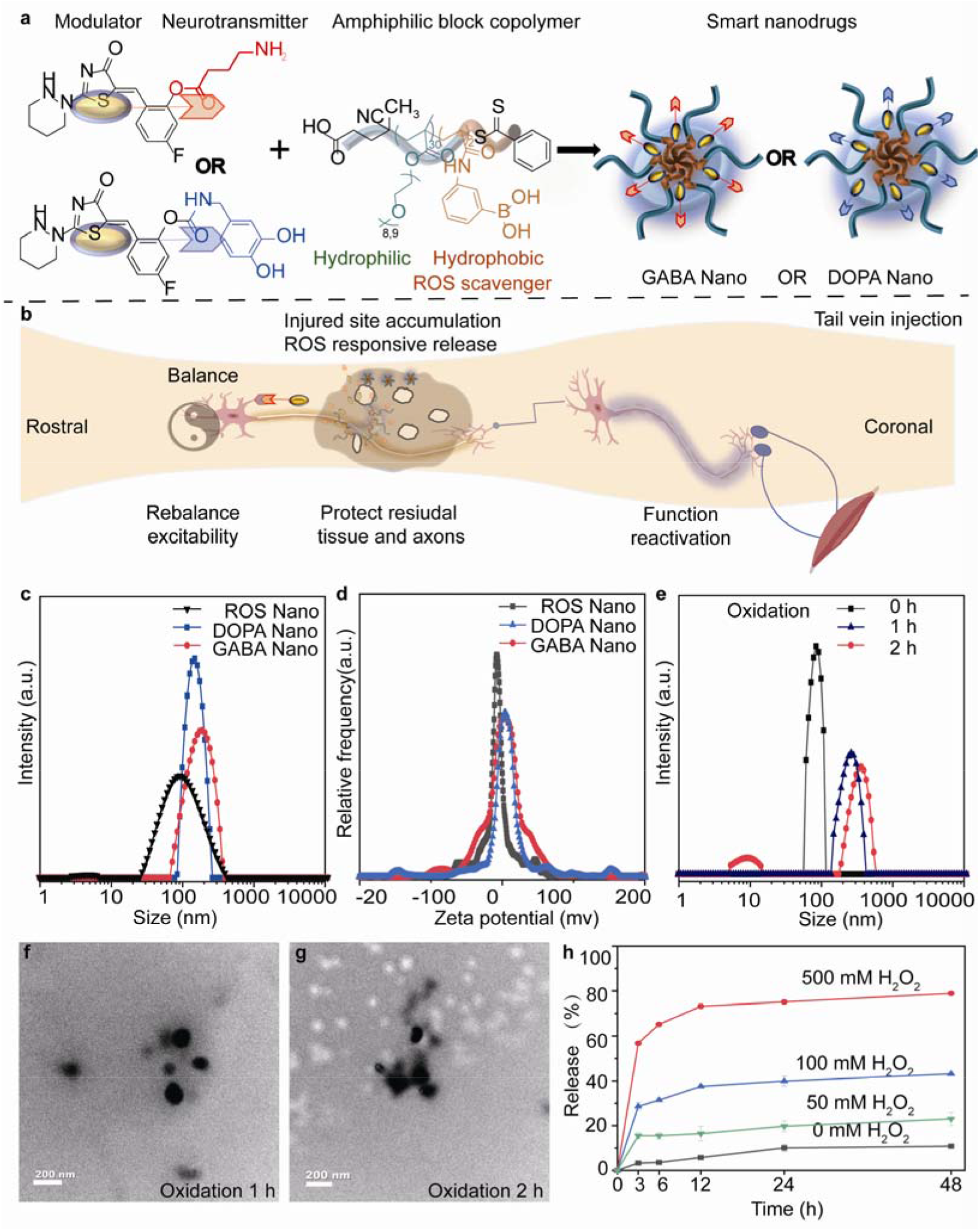
Design and characterization of the smart nanodrugs. **a,** Schematic depicting the synthesis of smart nanodrugs; nanoparticles share the same amphiphilic polymeric nanocarrier but have different neurotransmitter-functionalized modulators (GABA Nano or DOPA Nano). **b**, Schematic of the mechanisms of the smart nanodrugs after SCI. Following administration by tail vein injection, the smart nanodrugs accumulated in injury site through ROS-responsive release, consumed excessive ROS, and released neuron-targeted modulators. The effects of the nanodrugs include protecting spared tissue/axons, rebalancing the excitability of specific neurons, and eventually reactivating the lumbosacral central pattern generator (CPG) after SCI. **c-d**, The particle sizes and zeta potentials of empty nanoparticles (ROS Nano), DOPA Nano and GABA Nano. **e**, Changes in ROS nanoparticle size after treatment with H_2_O_2_ (500 mM). **f-g,** Representative TEM images of ROS Nano after oxidation by H_2_O_2_ for 1 h and 2 h. **h**, The release rates of ROS Nano in the presence of different concentrations of H_2_O_2_.

We applied this method to a clinically relevant severe contusive SCI model and demonstrated that the ROS-responsive nanodrugs protected spared axons from secondary injury. After undergoing ROS-responsive release, prodrugs from the nanodrugs expectedly accumulated at the injury site, successfully penetrated the BSCB and efficiently targeted the neurons of interest (Fig. 1b). Remarkably, the GABA, but not dopamine, nanodrug treatment promoted the integration of rescued dormant propriospinal circuits into local spinal cord circuitries and improved hindlimb locomotor function after SCI (Fig. 1b). This innovative pharmacological approach of modulating the activity of specific neurons provides an exciting direction for the treatment of SCI or other ROS-enriched neurological diseases.

## Result

### Synthesis and characterization of the designed nanodrugs

To specifically target the neurons of interest, the KCC2 agonist CLP-257^6, 21^ was conjugated to the neurotransmitter GABA or dopamine (Fig. 1a). The GABA- or dopamine-conjugated prodrugs release CLP-257 after hydrolyzation under physiological conditions. The structures of these synthesized prodrugs were confirmed by ^1^H-NMR (Fig. S1). However, the synthesized prodrugs were hydrophobic and had a poor solubility in physiological environments, limiting their applicability and efficacy for noninvasive administration. To overcome these limitations, an amphiphilic block copolymer with a polyethylene glycol (PEG)-based hydrophilic segment and a hydrophobic segment with boron-based ROS scavengers was designed to encapsulate these prodrugs. The block copolymer was synthesized through a reversible addition-fragmentation chain transfer polymerization of oligomer glycol monomethyl ether ester methacrylate (OEGMA) and 3-acrylamidophenylboronic acid (BAA) in an oxygen-free environment (Fig. S2a). ^1^H-NMR confirmed the structure of the synthesized copolymer (Fig. S2b-c). These block copolymers were expected to self-assemble into micelles when the concentrations of copolymers exceeded their critical micelle concentrations (CMC) in aqueous solutions.

Next, we sought to optimize the size of the self-assembled nanoparticles for optimal tissue penetration. As such, the copolymers were synthesized and characterized by varying the ratios of OEGMA:BAA monomers from 15:2 to 50:2 (Fig. S2d-e). The synthesized copolymers were then introduced into aqueous solutions to form micelles. Using a light scattering method, we found that the hydrodynamic diameters of the micelles ranged from 108 nm to 325 nm and their dispersity in aqueous solutions was excellent (PDI = 0.07-0.23). Because particles with an average size of 10-150 nm were reported to have better tissue penetration ability10, micelles with hydrodynamic sizes (ca. 108 nm) produced by the copolymer poly-2 were selected for further studies and were named ROS Nano (Fig. S2d-e). Further analysis with transmission electron microscopy (TEM) imaging showed that the self-assembled micelles were uniformly distributed with round morphologies. The average size of the nanoparticles was approximately 70 nm, which was slightly smaller than their hydrodynamic diameter (ca. 108 nm), and this suggests that dehydration occurred during the TEM preparation process (Fig. S2f).

Then, the prodrugs of CLP-257 conjugated with GABA or dopamine were encapsulated into the core of the ROS Nano through the nanoprecipitation method34. The fabricated nanodrugs were named GABA Nano and DOPA Nano, respectively. The characterization of these nanodrugs, including their critical micellar concentrations and drug loading capacity, is summarized in Fig. S2e. Both TEM images demonstrated round morphologies with an average size of approximately 90 nm (Fig. S2 g-h). The hydrodynamic diameters of the nanodrugs were slightly enlarged due to drug loading (Fig. 1c). Then, the surface potentials of these nanodrugs were characterized (Fig. 1d). The zeta potential of the empty nanodrug ROS Nano was approximately -10 mV before drug loading and changed to +1 and +5 mV after the dopamine-conjugated prodrug and GABA-conjugated prodrug were added, respectively, indicating that drug loading slightly changed the surface charge of the nanodrugs. Because of the inert and shielded PEG coating, the nanodrugs almost had a neutralized surface, which may prevent them from being recognized and ingested by macrophages before they reach the targeted site^35, 36^.

### The synthesized nanodrugs are responsive to ROS stimulation *in vitro*

We next attempted to determine whether the synthesized nanodrugs responded to ROS-enriched environments. To do this, we exposed the ROS Nano to H_2_O_2_ stimulation and analyzed its response with TEM and dynamic light scattering. We found that the ROS Nano responded quickly to H_2_O_2_ stimulation and showed noticeable expansion and collapse (Fig. 1e-g), which is consistent with the findings of previous studies^30, 32, 33^. Moreover, the release profiles of Nile Red encapsulated in ROS Nano were positively dependent on the H_2_O_2_ concentration (Fig. 1h), suggesting that the nanodrugs are highly sensitive to ROS stimulation. Due to this extraordinary feature, the nanodrugs were expected to respond to the ROS-enriched environments of SCI lesions.

Because cytotoxicity is crucial for in vivo studies, the cytocompatibility of the nanodrugs was also investigated. After performing a culture with the nanodrugs, we found that the morphologies of PC12 cells were normal even at the highest concentration of 500 μg/mL for 24 h, and the cell viability of PC12 was over 90% for all examined nanodrugs, indicating that these nanodrugs have a low cytotoxicity (Fig. S3a, b). To examine whether the nanodrugs could clear ROS and protect cells from oxidation damage, we induced PC12 cells to produce ROS with lipopolysaccharide (LPS) stimulation and then added nanodrugs to the culture medium. The results showed that the survival rate of cells treated with nanodrugs was remarkably higher than that of PBS-treated control cells, indicating that these nanodrugs have a good ability to protect cells against ROS damage (Fig. S3c). Furthermore, compared with cells that received the PBS treatment, the ROS level was obviously decreased in the cells that received the nanodrug treatment, verifying that these nanodrugs have a good ability to scavenge ROS (Fig. S3d, e). Altogether, the designed nanodrugs exhibit perfect cytocompatibility and excellent antioxidative performance, indicating that they have a reliable ability to protect cells against ROS damage.

### Designed nanodrugs exhibit intensive spinal cord lesion accumulation and neuron-targeting properties

SCI damages the microvasculature of the spinal cord and leads to acute breakdown of the BSCB. Subsequently, the recruited inflammatory cells accumulate in the lesion site and generate an environment with extensive amounts of toxic ROS31. Since the nanodrugs intensively responded to ROS stimulation, we reasoned that they could responsively release their payloads of prodrugs in the ROS-enriched environment of the injured spinal cord (Fig. 2a). To test this hypothesis, we first optimized a severe tenth thoracic (T10) contusive SCI model, in which the contusion was prolonged to 5 seconds to maintain a consistency injury severity. In this study, we replaced CLP-257 with a fluorescence dye, Cy5.5, to investigate the drug release profiles of GABA Nano and DOPA Nano. The near-IR dye Cy5.5 was chosen to avoid an overlap of signals with the strong autofluorescence from blood cells in the lesion. SCI rats were randomly administered 10 mg/kg Cy5.5-conjugated GABA Nano (GABA Nano@Cy5.5) or DOPA Nano (DOPA Nano@Cy5.5) through an injection in their tail veins 3 h after SCI, and this represented the earliest clinically feasible time point for such a treatment. The spinal cords of the rats were collected at 3, 6, 24 and 48 h after nanodrug administration for further analysis (Fig. 2b).

**Fig. 2.**
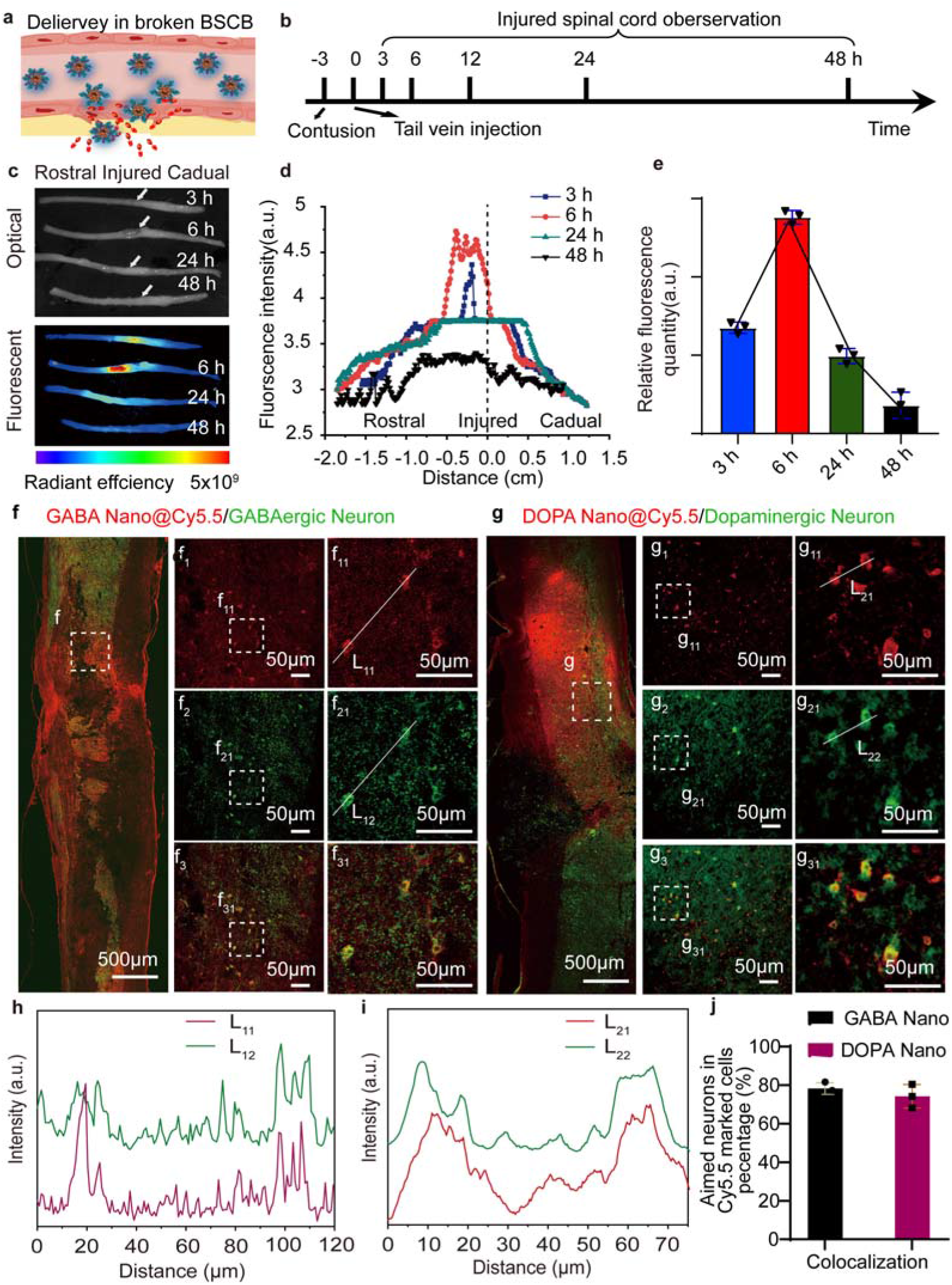
Smart nanodrugs accumulate at the injury site in the spinal cord and selectively target specific neurons. **a**, Schematic of the nanodrugs reaching and entering the spinal cord following BSCB disruption. **b**, Experimental schedule of SCI, drug delivery and observation. **c,** Images of spinal cords at the indicated time points after SCI and injection of ROS Nano@Cy5.5. **d-e**, Spatial and temporal quantification of the fluorescent intensity of ROS Nano@Cy5.5 in the injured spinal cord. **f**, Representative longitudinal images of tissues from the GABA Nano@Cy5.5 (red) injection group stained with anti-GABA antibody (green). Regions in the white squares (f_1_-f_3_ and f_11_-f_31_) are enlarged on the right side and show details of GABA Nano-marked cells and GABAergic neurons near the injury sites. **g**, Representative longitudinal images of tissues from the DOPA Nano@Cy5.5 (red) injection group stained with anti-DOPA antibody (green). Regions in the white squares (g_1_-g_3_ and g_11_-g_31_) are enlarged on the right side. **h-i**, Intensity profiles of the white lines in Fig. 2f and 2g. **j**, Quantitative analysis of targeting efficiency via evaluation of the colocalization percentage of target neurons in Cy5.5 marked cells. n = 3 for each group. Data are shown as mean ± SEM.

We found that the nanodrugs intensively accumulated around the spinal lesion site 3 h after injection, reached the maximum concentration at approximately 6 h after injection, and persisted for 48 h. This indicates that sufficient spinal tissue had accumulated and that the nanodrugs had a long circulation time, which might benefit in vivo targeted delivery (Fig. 2c). Interestingly, fluorescence distribution analysis indicated that the nanodrugs preferentially accumulated in the rostral region adjacent to the lesion, possibly due to the occurrence of spinal vascular occlusion after severe SCI (Fig. 2d)^37, 38^. The tropism of these nanodrugs in this SCI model facilitates the targeting of propriospinal neurons above the lesion with spared descending projections.

To further characterize the tropism of nanodrugs in the injured spinal cord, longitudinal spinal cord sections were immunostained with GABAergic and dopaminergic neuron markers and were subjected to colocalization analysis. We found that a number of GABA Nano labeled cells (red) were colocalized with cells that were stained with GABAergic markers (green) (Fig. 2f and h). Similarly, numerous DOPA Nano-marked cells (red) were colocalized with cells that were stained with dopaminergic markers (green) (Fig. 2g and i). Through quantitative analysis of different rats, we found that approximately 80% of GABA Nano- or DOPA Nano-marked cells were their target neurons (Fig. 2j). These results indicated that these nanodrugs have a high selectivity and efficiency.

To investigate the biodistribution of these nanodrugs, the NIR dye Cy5.5 was encapsulated into the nanodrug delivery system (ROS Nano@Cy5.5), facilitating the tropism assessment of these nanodrugs through fluorescent imaging. The main organs in the rats injected with ROS Nano@Cy5.5 were dissected and analyzed by fluorescence imaging. We found that most of the nanodrugs were present in the liver and kidney in the first 3 h, as expected, while the concentration of nanodrugs in the kidney exceeded that in the liver 6 h after injection, suggesting that kidney-mediated excretion of these nanodrugs had occurred (Fig. S4a, b). In addition, the histological sections of the heart, kidney, liver, lung, and spleen of the rats with nanodrugs or PBS injections showed no significant differences, and this suggests that these nanodrugs have a low toxicity (Fig. S4c). Taken together, the results indicate that the designed ROS-responsive nanodrugs have low toxicity and competitively target neurons of interest in the injured spinal cord.

### The nanodrug delivery system could deliver small molecules to the spinal cord when the BSCB is restored

It is known that the BSCB is restored within 7 days post-SCI9, and this prevents macromolecular agent from penetrating in the late stage of SCI39. Fortunately, the restored BSCB still allows the penetration of small molecules or specific peptides or proteins40. Because SCI-induced ROS upregulation lasts for months41, we speculated that the designed nanodrugs might release the encapsulated prodrugs in response to ROS, which could in turn penetrate the injured spinal cord (Fig. 3a). To test this hypothesis, in the same SCI model, we intravenously administered 10 mg/kg nanodrug of ROS Nano@Cy5.5 to different groups of rats, including intact rats and rats at 4, 7, and 14 days after SCI (Fig. 3b). The rats were sacrificed 6 h after the nanodrug injection for analysis. As a result, we found that the nanodrug could transport Cy5.5 into the spinal cord adjacent to the lesions in the rats with injection at 4 days or later after SCI (Fig. 3b-c). Surprisingly, the fluorescence intensity was much stronger in the rats with injection at 7 days post-SCI than in the rats with injection at 4 or 14 days post-SCI (Fig. 3b-c), which is contradictory to our expectation because the BSCB was more refractory to penetration in the late stage of SCI. A rational explanation for this observation is that drug penetration is not dependent on the penetration of nanoparticles but rather on the disassembly efficiency of nanoparticles. As the ROS level reached its peak at 7 days post-SCI42, the nanodrugs could disassemble more efficiently and release their small molecules cargo. Ultimately, more small-molecule compounds penetrated the restored BSCB (Fig. 3b-c). However, almost no fluorescence was detected in the spinal cords of intact rats, further confirming that small molecule penetration in the BSCB was dependent on ROS stimulation (Fig. 3d). To further validate this BSCB penetration, longitudinal spinal cord sections from these rats were analyzed (Fig. 3e). We found that most Cy5.5-marked cells (magenta) were colocalized with neurons (red) but not GFAP-marked astrocytes (green), suggesting that Cy5.5 penetrated the BSCB and preferentially marked neurons. In addition, a higher number of Cy5.5-marked cells (magenta) was observed in the rats that were with injection at 7 days post-SCI, and almost no Cy5.5-marked cells were observed in intact rats, which was consistent with previous observations. Thus, it is conclusive that the drug delivery system is able to deliver hydrophobic small-molecule compounds into the spinal cord even when the BSCB is restored in the late stage of SCI.

**Fig. 3.**
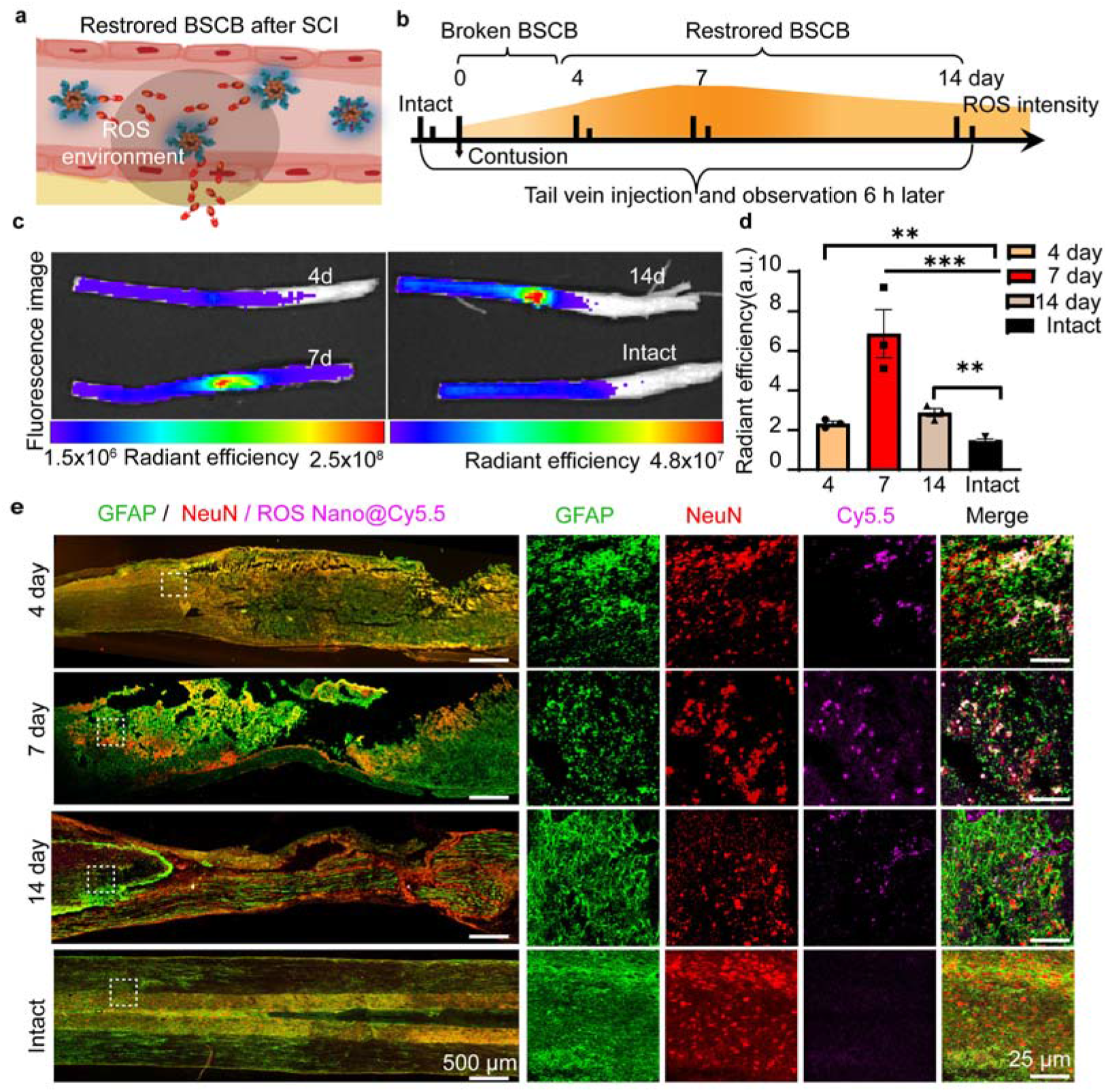
The ROS-responsive nanodrug delivery system delivers small molecules into the spinal cord even after BSCB restoration. **a**, Schematic depicting the penetration method of the nanodrug delivery system following restoration of the BSCB. **b**, Schematic diagram of the experimental design. **c**, Representative fluorescent images of spinal cords 6 h after injection. ROS Nano@Cy5.5 was injected at 4, 7 or 14 days post-SCI. **d**, Quantitative analysis of spinal cords following ROS Nano@Cy5.5 injection. Data are shown as mean ± SEM. One-way ANOVA with Tukey’s post hoc test was used for comparisons among multiple groups. n = 3. **P < 0.01, ***P < 0.001 statistical significance. **e**, Representative images of spinal cord sections stained with anti-GFAP antibody (green) and anti-NeuN antibody (red) in ROS Nano@Cy5.5 (magenta)-injected spinal cords.

### Nanodrugs protect spared tissues/axons from secondary injury

After verifying that our nanodrugs were capable of scavenging ROS in vitro and delivering drugs to target neurons, we attempted to assess whether this treatment could ameliorate the SCI-induced neuroinflammatory environment and protect spared tissues/axons from secondary injuries. To this end, we employed the same severe T10 contusive SCI model as that used in the drug release study. Unlike the moderate contusive SCI model with large individual variations, the rats with severe contusive SCI showed minimum ankle movement and exhibited nearly complete hindlimb paralysis within 10 weeks after injury. In this study, the examiners were blinded to the drug treatments and 10 mg/kg ROS Nano or PBS was intravenously administered to rats at serial time points after SCI, as detailed in Fig. 4a. In the ROS Nano or PBS treatment groups, several rats terminated on the 7^th^ day after SCI. The spinal cords were dissected for a western blot analysis to evaluate the protein expression level in the spinal lesions of rats with different treatments. Rat hindlimb locomotion was recorded by a camera for later behavioral analysis. Finally, the rats with 9-week behavioral recordings were sacrificed for a histological study.

**Fig. 4.**
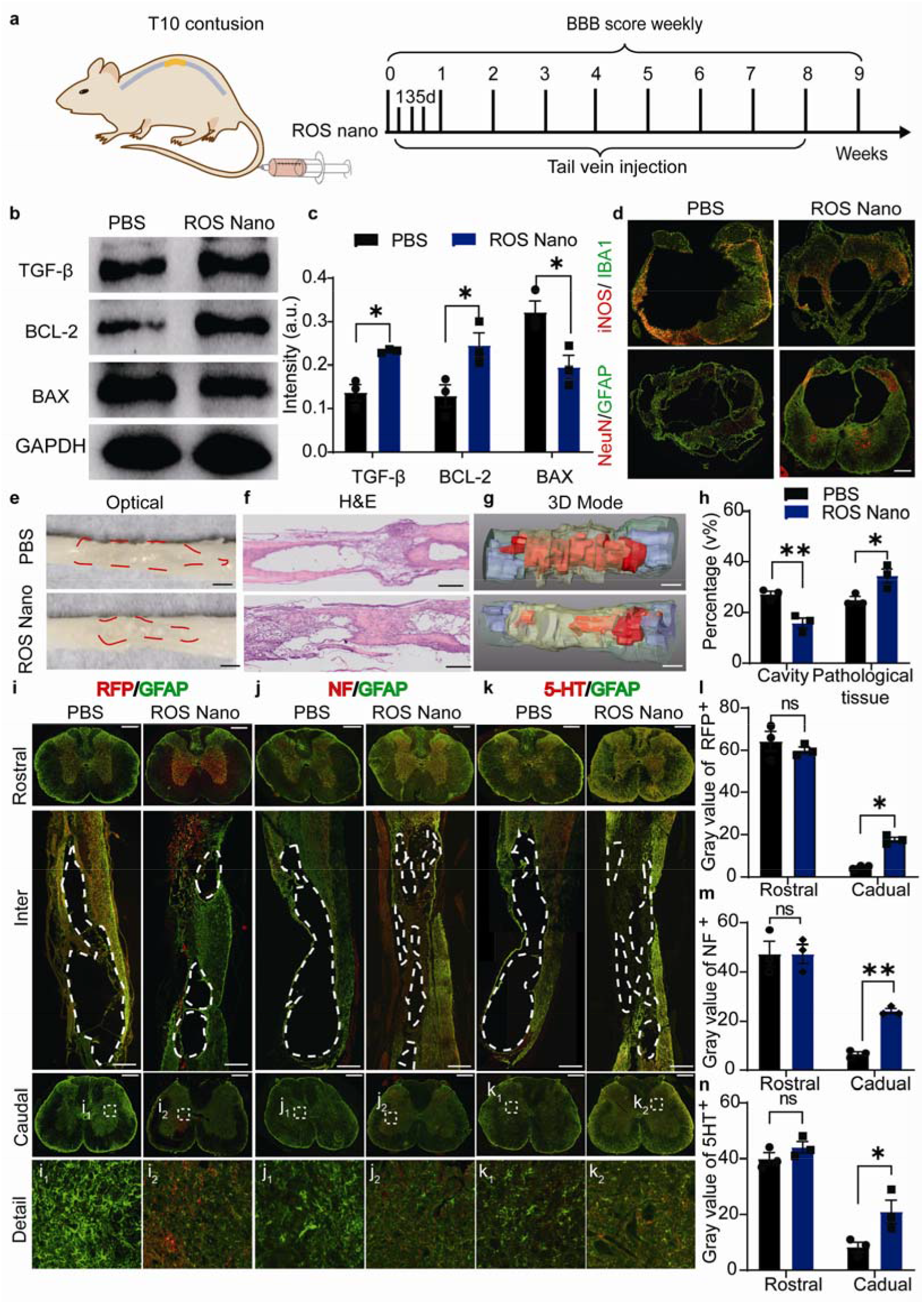
The ROS Nano treatment protects spared tissues/axons from secondary injury. **a**, Schematic diagram of the experimental design. **b-c,** Representative immunoblots and quantitative analyses of TGF-β, BCL-2 and BAX expression in the injury site 1 week after PBS or ROS Nano (10 mg/kg) treatment. n = 3. * indicates p < 0.05. Data are shown as mean ± SEM. **d**, Representative images of transverse sections at the epicenter of an injury stained with iNOS (red)/IBA1 (green) or NeuN (red)/GFAP (green) after 4 weeks of treatment. **e-f**, Representative optical images and H&E-stained images of longitudinal sections of injured spinal cords after 9-weeks treatment. The boundaries of the injury sites are marked with red dotted lines. Scale bars, 1 cm (e) and 1 mm (f). **g**, 3D reconstruction images of spinal cord tissues showing white matter (green), gray matter (light blue), pathological tissue (yellow), and cavities (red). Scale bar, 1 mm. **h**, Graph showing the quantification of the cavity and pathological volume percentages at the injury site. Data are shown as mean ± SEM. One-way ANOVA with Tukey’s post hoc test was used for comparisons among multiple groups, and two-tailed paired t tests were used for comparisons between two groups. n = 3. * and ** indicate p < 0.05 and p < 0.01, respectively. **i-k**, Representative images of spinal cord sections stained with GFAP (green), NF (red), 5-HT (red) or RFP (red) of rats after 9 weeks of treatment with PBS or ROS Nano. The white dotted lines indicate the boundaries of the cavities. Rostral (upper), longitudinal (middle) and caudal (lower) spinal cord sections. The details are shown in the enlarged images of i_1_, i_2_, j_1_, j_2_, k_1_, and k_2_. **l-n**, Graph showing the quantification of the RFP, NF and 5-HT immunoreactivity gray values on the rostral and caudal sides. Data are shown as mean ± SEM. One-way ANOVA with Tukey’s post hoc test was used for comparisons among multiple groups, and two-tailed paired t tests were used for comparisons between two groups. n = 3. *and ** indicate p < 0.05 and p < 0.01, respectively.

We found that the expression of the anti-inflammatory marker TGF-β and the antiapoptotic marker Bcl-2 in the lesion sites was remarkably increased with the ROS Nano treatment (Fig. 4b-c). In contrast, the proapoptotic marker BAX was significantly reduced, suggesting that the ROS Nano treatment ameliorated the SCI-induced neuroinflammatory environment in the lesion. In addition, copolymer-based micelles might also help seal the cell membrane and reduce damage from the influx of Ca^2+^ 43. To further analyze the tissue-protective efficacy, epicenter cross sections of injured spinal cords at 4 weeks post-SCI were analyzed by immunofluorescence staining (Fig 4d). Section images with IBA1 (macrophages/microglia) and iNOS (activated immune cells) staining showed that the activated immune cells were significantly reduced through the ROS Nano treatment compared with that of the PBS control. More neurons (NeuN^+^) and astrocytes (GFAP^+^) survived in the rats treated with ROS Nano (Fig 4d), suggesting that this treatment protected spared tissues from secondary injury in SCI lesions.

Furthermore, we found that the shape deformation of the spinal cord was obviously less in the rats injected with ROS Nano 9 weeks post-SCI than in the PBS control group (Fig. 4e), and the hematoxylin-eosin (H&E)-stained images of the spinal cord sections showed that the cystic cavity in the ROS Nano treatment group was notably smaller than that in the PBS-treated rats, which extended rostrocaudally more than 5 mm away from the epicenter of the injury site (Fig. 4f). To quantitatively analyze the cavity volume, images of spinal cross sections were collected in sequence and reconstructed to 3D models (Fig. 4g). After quantification across different rats, we found that in the PBS-treated rats, approximately 27.3% of the spinal cord volume was cavities. In contrast, the cavity volume percentage in the ROS Nano-treated rats was significantly decreased to approximately half of that (approximately 15.8%) in the PBS-treated rats (Fig. 4h). Compared to the PBS treatment, tissue with abnormal architecture, which was defined as pathological tissue, was increased in the ROS Nano rats at the injured site (ca. 35% vs. 25%).

We next attempted to determine whether more descending axons could survive with this treatment. To this end, AAV2/9-mCherry was injected into T7-T8 of some rats 2 weeks before they were sacrifices to trace the descending propriospinal axons, and immunostaining of red fluorescent protein (RFP), anti-neurofilament (NF) and anti-5-hydroxy tryptamine (5-HT) was performed to visualize descending propriospinal axons, ubiquitous axonal fibers, and serotonergic axons, respectively (Fig. 4i-n). We found that very few axons could extend into the lesion site in all examined subjects. In the coronal sections of spinal segments, RFP^+^ propriospinal axons, ubiquitous axonal fibers, and serotonergic axons caudal to the lesion were remarkably higher in the rats treated with ROS Nano than in the PBS-treated rats (Fig. 4i-n). Serial images of coronal spinal cord sections showed that there were more spared RFP^+^ propriospinal axons in the ROS Nano-treated spinal cord than in the PBS-treated spinal cord, and these axons were located around the central injured site and extended into the caudal spinal cord (Fig. S5). These results suggested that the axons observed below the lesion sites were spared axons that were recused from secondary injury.

### The rescued spared axonal connections mediate limited hindlimb locomotor recovery

After verifying that the spared axons could be rescued efficiently through ROS Nano treatment, we next attempted to determine whether these axons could mediate hindlimb locomotor functions. After blinding the identity of each rat group to the examiners, the previously mentioned videos of the hindlimb performances of rats were analyzed with BBB scoring. As shown in Fig. S6a, severe SCI caused complete hindlimb paralysis in all examined rats at 1 week postinjury (BBB score of 0). Nine weeks after SCI, the rats treated with PBS (n = 10) showed limited spontaneous recovery that plateaued at 4 weeks postinjury, with an average BBB score of 0.5 (only isolated ankle movement). This is possibly because almost no descending input extended into the lumbosacral CPG and mediated hindlimb locomotion in this condition. In contrast, the BBB scores of rats treated with ROS Nano slowly increased to approximately 2 in 4 of the 10 rats, and these rats performed vigorous ankle movements; however, their limbs did not support their weight and they did not have an ability to step. The joints, angle movements and height aptitudes involved in the ability of rats to walk are shown in Fig. S6b, and the representative hindlimb kinematics are shown in Fig. S6c and 6d. Interestingly, some of the rats in the ROS Nano group showed muscle spasm-like episodes, which was also indicated by the unregulated tibialis anterior (TA) muscle activity in the MEG recordings of these rats (Fig. S6e). In conclusion, weekly ROS Nano treatment resulted in some effects that improved hindlimb locomotion; however, these effects lacked statistical significance (P = 0.16). These results indicate that the spared descending axons that were rescued were unable to mediate significant hindlimb locomotion functional recovery in the disordered spinal network after severe SCI.

### GABA Nano treatment improves hindlimb locomotor recovery after SCI

We speculated that the reason why propriospinal neurons with spared descending axons could not mediate hindlimb locomotion is that they were hyperactive after SCI and thus unable to efficiently relay the signals from the brain to the lumbosacral CPG. Therefore, we sought to assess whether the neuron-targeting treatment would promote the integration of spared descending axons into the host spinal network and perform hindlimb locomotor function. Through histological analysis, we found that the GABA Nano and DOPA Nano treatment showed similar spared tissue/axon protective effects as ROS Nano (Fig. S7). Interestingly, most of the rats treated with DOPA Nano only displayed ankle movements with a plateaued BBB score of approximately 2 at the test time points (Fig. 5a, b), whereas rats treated with GABA Nano exhibited significantly improved hindlimb locomotor functions, performing hindlimb plantar placement or dorsal stepping at 5 weeks postinjury (P<0.01) (Fig. 5a, b). More strikingly, 3 of the 10 rats that were treated with GABA nano recovered the ability to consistently take hindpaw plantar steps with occasional weight support (BBB > 8, Fig. 5b, Supplementary Movie S1). These results suggested that the GABA Nano treatment could markedly improve hindlimb motor function and weight-bearing stepping after severe SCI, and this was implicated as the limiting step for functional recovery in such a severe SCI model44. Detailed hindlimb kinematics revealed the following significant improvement with this GABA Nano treatment: (1) increased weight support (increased iliac crest height, Fig. 5d, h, S8a); (2) increased max toe high and high aptitude (Fig. S8c, d), and (3) strikingly increased ankle amplitude (Fig. 5d, h, S8e-g).The variety of functional recovery with this treatment is possibly because the rescued descending axons that are closely relevant to hindlimb locomotion are randomly presented in the injured rats. Moreover, the BBB scores were maintained for 2 weeks after the treatment was stopped (Fig. S9), suggesting that the sustained functional recovery was due to the GABA Nano treatment. In a complete crush injury model, we found that the GABA Nano treatment could not promote significant functional recovery (Fig. S10), suggesting that hindlimb functional recovery with this treatment relied on spared spinal connections across the lesion.

**Fig. 5.**
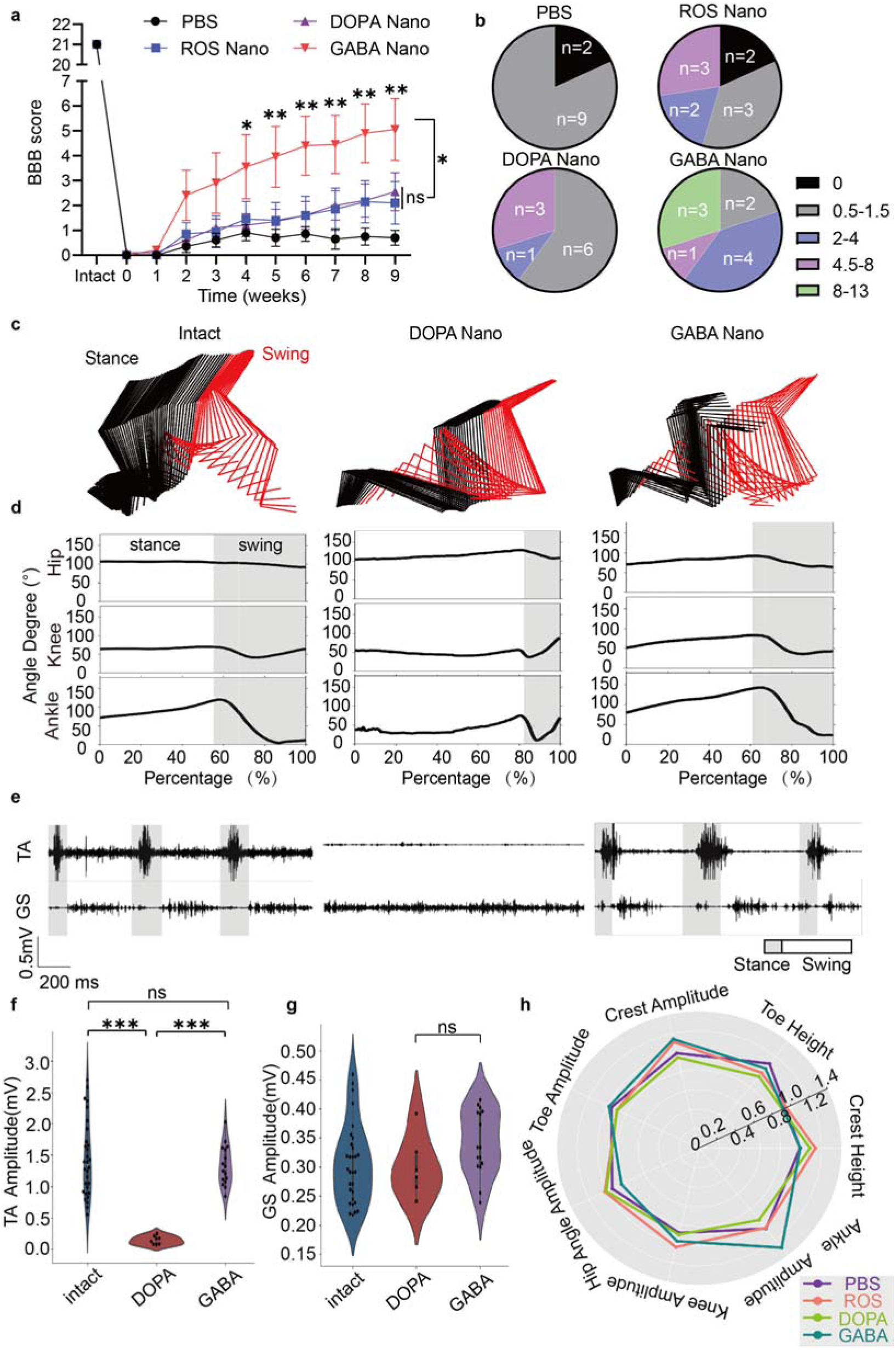
GABA Nano treatment improves the recovery of hindlimb locomotor functional in rats with severe contusive SCI. **a**, Weekly BBB scores of the rats treated with PBS, ROS Nano (10 mg/kg), DOPA Nano (10 mg/kg) or GABA Nano (10 mg/kg). Data are shown as mean ± SEM. One-way ANOVA with Tukey’s post hoc test was used for comparisons among multiple groups, and two-tailed paired t tests was used for comparisons between two groups. n = 10. *and ** indicate p < 0.05 and p < 0.01, respectively. **b**, Bar graphs showing the distribution of BBB scores for rats with PBS, ROS Nano, DOPA Nano or GABA Nano treatments. **c**, Representative color-coded stick views of kinematic hindlimb movement of the intact rats or rats treated with DOPA nano or ROS Nano. **d**, Representative curves of the hip, knee and ankle angles during a one-step cycle. **e**, Representative EMG data for the TA and GS muscles from the intact, DOPA Nano and GABA Nano groups. **f-g**, Quantitative analysis of signal amplitudes from TA and GS muscles for rats with different treatments. Data are shown as mean ± SEM. One-way ANOVA with Tukey’s post hoc test was used for comparisons among multiple groups, and two-tailed paired t tests were used for comparisons between two groups. n = 5 to 12. *and ** indicate p < 0.05 and p < 0.01, respectively. **h**, Well-rounded quantification of seven described behavioral features in animals subjected to the indicated treatments through a radar graph.

To analyze the muscle activity of these rats, we recorded the electromyogram (EMG) of hindlimb muscles. We found that the ankle flexor TA muscle and extensor gastrocnemius soleus (GS) muscle of rats treated with ROS Nano or DOPA Nano showed some activities during hindlimb joint movement, while they were rarely active in rats treated with PBS (Fig. 5e). Interestingly, while the rhythm of the GS and TA muscles was compromised in rats treated with GABA Nano compared with intact rats, the GS muscles in rats treated with DOPA Nano showed a high amplitude without any rhythm, which explains why they did not exhibit a normal gait (Fig. 5e, S8h, i).

### GABA Nano treatment alters the excitability of interneurons and reanimates lumbosacral CPG

It is known that reducing the excitability of inhibitory interneurons after SCI facilitates functional restoration12. To assess whether GABA Nano treatment improved functional recovery through a similar mechanism, we used c-Fos immunoreactivity as a proxy to detect the neuronal activity of the interneurons in the spinal cord. After walking on a treadmill for 1 h, the rats were sacrificed, and their T8 and L2 spinal cord segments were collected and stained with c-Fos and NeuN (a neuronal marker) at 9 weeks after SCI. As shown in Fig. S11a and b, the c-Fos-positive cells in the spinal segments were largely costained with NeuN, suggesting that c-Fos immunostaining has a high specificity for neurons. Representative composites of c-Fos/NeuN double-positive cells are illustrated in Fig. 6a. While the c-Fos^+^ neurons of intact rats were distributed homogenously in T8 and L2 spinal segments, the c-Fos^+^ neurons in SCI rats with the PBS or ROS Nano treatment were concentrated in the dorsal horn of the T8 spinal cord and they rarely presented in the L2 spinal cord below the lesion, which was consistent with the observation in other SCI models45 (Fig. 6a-d, S11). However, with the GABA Nano treatment, the intensive distribution of c-Fos^+^ neurons in the dorsal horn of the T8 spinal cord was remarkably reduced, and more c-Fos^+^ neurons were present in the intermediate and ventral spinal cord, indicating that they developed a distribution pattern similar to what was detected in intact rats (Fig. 6a-d, S11). On the other hand, a significant increase in the number of c-Fos^+^ neurons was detected in the L2 spinal cord of rats treated with GABA Nano compared with the PBS-treated control rats (Fig. 6a-d, S11). The distribution of c-Fos^+^ neurons in the T8 spinal cord was slightly changed in the DOPA Nano-treated rats compared with the GABA Nano-treated rats (Fig. 6a-d, S11). This may have been because dopaminergic neurons in the spinal cord are mainly responsible for physiological functions other than motor control46. Conclusively, these findings suggest that the GABA Nano targeting treatment transforms the SCI-induced irregulated activity pattern of interneurons into a more physiological state and allows the spared propriospinal connections to integrate into local spinal circuitry and re-engage lumbosacral CPG to perform functions (Fig. 6e).

**Fig. 6.**
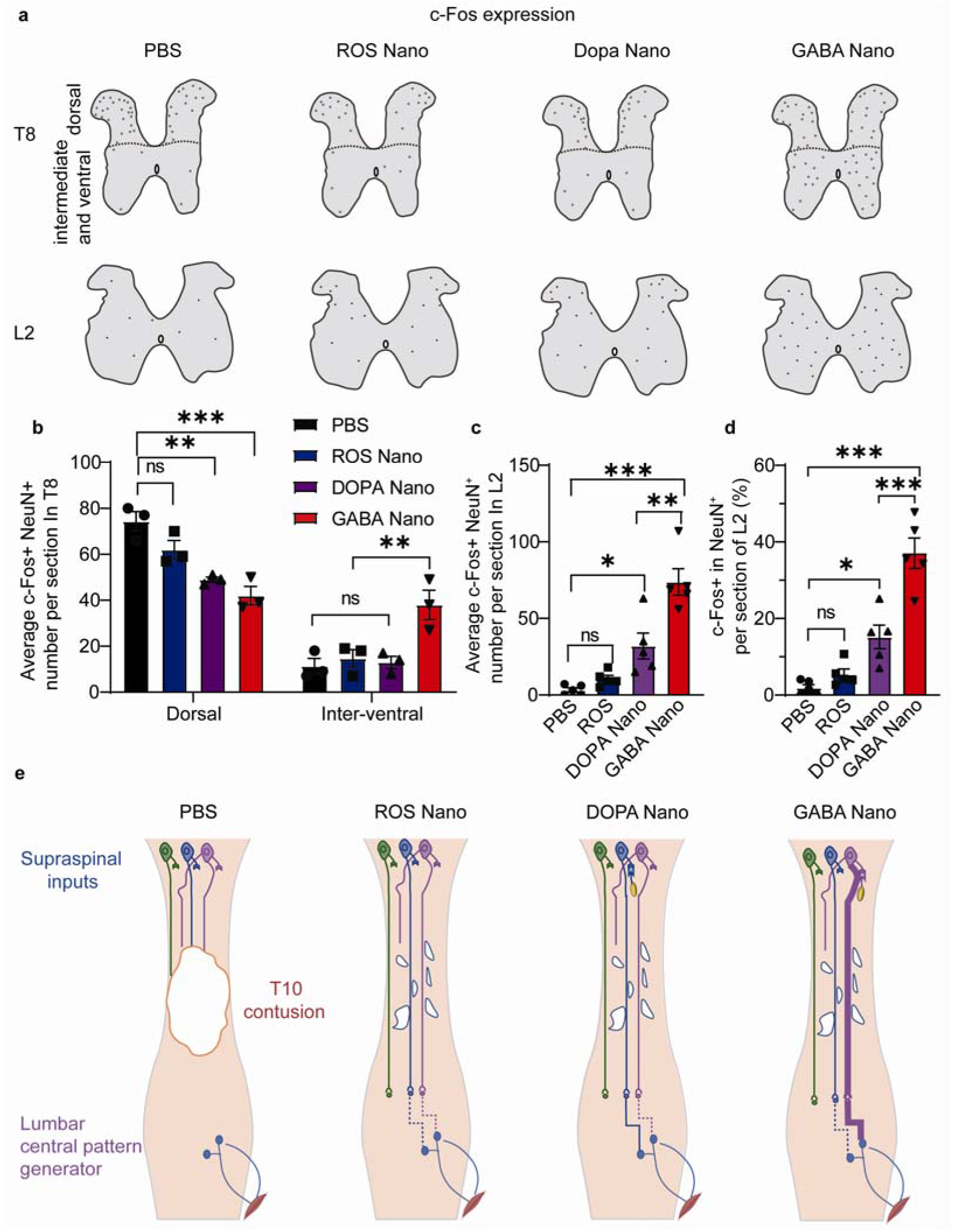
GABA Nano treatment rebalances neuronal activity and activates rescued residual spinal circuits to perform functions. **a**, Schematics of spinal cord cross sections showing c-Fos expression (representing neuronal activity) patterns in T8 and L2 segments after 1 h of continuous locomotion of injured rats treated with PBS, ROS Nano, DOPA Nano or GABA Nano. Each spot represents a cell positively stained with both c-Fos and NeuN. Representative raw images are shown in fig S11. **b**, Quantification of c-Fos^+^ neurons in dorsal or in interventral of T8 sections. Data are shown as mean ± SEM. One-way ANOVA with Tukey’s post hoc test was used for comparisons among multiple groups, and two-tailed paired t tests were used for comparisons between two groups. n = 3. *and ** indicate p < 0.05 and p < 0.01, respectively. **c**, Quantification of c-Fos^+^ neurons in L2 sections of all groups. Data are shown as mean ± SEM. One-way ANOVA with Tukey’s post hoc test was used for comparisons among multiple groups, and two-tailed paired t tests were used for comparisons between two groups. n = 5. *and ** indicate p < 0.05 and p < 0.01, respectively. **d**, Quantification of the c-Fos^+^ percentage in NeuN^+^ cells in L2 sections of all groups. Data are shown as mean ± SEM. One-way ANOVA with Tukey’s post hoc test was used for comparisons among multiple groups, and two-tailed paired t tests were used for comparisons between two groups. n = 5. *and ** indicate p < 0.05 and p < 0.01, respectively. **e**, Schematic of the effects of different treatments on the spinal cord after T10 contusion.

## Discussion

Here, we described here a noninvasive pharmaceutical nanodrug that both protects spared propriospinal connections from secondary injury by reducing ROS levels and transforms rescued dysfunctional spinal connections into a functional state after severe SCI. The inhibitory neuron-targeting treatment reduced the excitability of propriospinal neurons with remnant spinal connections above the injury site and thus facilitated a relay of the brain-derived commands to the lumbar spinal cord and reanimated lumbosacral CPG to perform functions. Our findings pave the way for the development of a noninvasive pharmacological treatment for severe SCI.

This study provides a novel strategy for spinal cord drug delivery in SCI treatment. It is known that nanodrugs can passively leak into acute spinal cord lesions following systemic administration. However, restoration of the BSCB in the chronic phase of SCI prevents nanodrugs from being consistently delivered. Traditionally, nanoparticle surfaces are modified with BSCB penetration peptides/proteins to enhance the BSCB penetration capacity of nanoparticles10. However, it is difficult for nanoparticles to directly penetrate the BSCB due to their size. Unlike the penetration strategies proposed by previous studies, the ROS-responsive spinal drug delivery system described by this study does not require the nanoparticles to penetrate the BSCB. Taking advantage of the persistent ROS-enriched environment of spinal cord lesions, the nanodrugs responsively released the prodrug, which intensively accumulated around the lesion and efficiently penetrated into the spinal cord in the end. Thus, ROS stimulus-responsive nanodrugs enhance targeted delivery and avoid undesirable exposure, further improving efficacy.

This study provides a novel pharmacological approach to target specific subtypes of neurons to treat SCI. Our approach achieved a remarkably high efficiency of approximately 80% toward the target neurons. Thus, the approach represents a novel strategy for targeting specified neurons for precise treatment. Among the different targeted treatments, we found that the inhibitory neuron-targeting approach is vital for functional recovery because the nonspecific neuron (DOPA)- and excitatory neuron-targeting methods were ineffective (data not shown). In addition, through targeted administration, the dosage required for treatment could be significantly reduced, and side effects could be eliminated by minimizing systemic distribution. In addition to treating SCI, this study opens a new exciting perspective for the treatment of other neurological diseases, such as neuropathic pain, absence seizures, SCI-induced spasticity and amyotrophic lateral sclerosis, that are caused by excitation/inhibition imbalance in specific regions^18, 20, 47, 48^.

In contrast to previous studies that focused on relay or endogenous remnant spinal circuitries^6, 9^, this study aimed to reactivate the dormant propriospinal circuits that were rescued from secondary injury. We found that the hindlimb locomotion restoration promoted by the GABA Nano treatment relied on spared propriospinal connections across the injury site. In a complete SCI injury model without spared spinal connections, the nanodrug treatment was not effective in functional restoration, possibly because the pharmacological treatment was not efficient enough to activate lumbosacral CPG through endogenous propriospinal circuits below the injury. Despite this, the results indicated that fine-tuning the excitability of propriospinal neurons with spared connections was a feasible method for restoring lost locomotor functions.

Even though the drug CLP257 has been proven to be able to penetrate BSCB6, the drug delivery efficiency of this system has to be accurately determined before clinical studies can be initiated. Moreover, no side effects were caused by this nanodrug, and systemic toxicity was not detected in our study; however, the long-term effects of nanodrug administration remain to be clarified. Although this study verified the treatment efficacy of nanodrug administration, the optimal dosage, administration frequency and timing of the treatment need to be optimized in future studies to achieve the best clinical outcomes. Exercise contributes to the restoration of endogenous inhibition49; thus, future studies may need to assess whether combining this nanodrug treatment with rehabilitation training could further improve its therapeutic effects.

## Method

### Animals and materials

All animal protocols were approved by the Institutional Animal Care and used following the provisions of Zhejiang University Animal Experimentation Committee (ZJU202010110). Sprague–Dawley rats (200-250 g, female) were purchased from the Experimental Animal Center of the Zhejiang Academy of Medical Science, Hangzhou, China. Chain transfer agent (CTA): 4-cyano-4-(phenylcarbonothioylthio) pentanoic acid; monomers: poly (ethylene glycol) methyl ether methacrylate (OEGMA, Mn=475) and 3-acrylamidophenylboronic acid (BAA); and initiator: 2,2’-azobis(2-methylpropionitrile) (AIBN) were purchased from Aladdin Reagent Co., Ltd. Monomers and initiators were purified by either recrystallization or filtration through an Al_2_O_3_ column to remove inhibitors. Dopamine hydrochloride (DOPA), γ-aminobutyric acid (GABA), N, N’-carbonyldiimidazole (CDI), fluorescein 5 (6)-isothiocyanate (FITC), and Cy5.5 NHS ester were purchased from Sigma (US). Dialysis bags, phosphate-buffered saline (PBS), Cell Counting Kit-8 (CCK-8) and 2′,7′-dichlorofluorescin diacetate (DCFH-DA) were obtained from Shanghai Yuanye Biological Technology Co., Ltd. Hydrogen peroxide (H_2_O_2_), dimethyl sulfoxide (DMSO), tetrahydrofuran (THF), N,N-dimethylformamide diethyl acetal (DMF), and diethyl ether were also purchased from Aladdin Reagent (Shanghai) Co., Ltd. The KCC2 agonist (Z)-5-(4-fluoro-2-hydroxybenzylidene)-2-(tetrahydropyridazin-1(2H)-yl)thiazol-4(5H)-one (CLP-257) was purchased from DC chemicals Co., Ltd. Methanol-d_4_, chloroform-d, and dimethyl sulfoxide-d_5_ were also purchased from Aladdin Reagent Co., Ltd. RPMI 1640 medium, fetal bovine serum (FBS), horse serum, and 1% penicillin/streptomycin were purchased from Gibco, and l-glutamine was obtained from Adamas. A live/dead viability/cytotoxicity kit was purchased from Thermo Fisher Scientific (China) Co., Ltd. A BCA protein assay kit and ECL kit were provided by Solabo Co., Ltd. (Shanghai, China).

Chicken anti-GFAP [Abcam (ab134436), 1:500], rabbit anti-neurofilament (NF) heavy polypeptide [Abcam (ab8135), 1:500], goat anti-5-HT antibody [Invitrogen (pa1-36157), 1:500], rabbit anti-iNOS [Abcam (ab283655), 1:200], goat anti-IBA1 [Abcam (ab5076), 1:500], Rabbit anti-NeuN [Abcam (ab177487), 1:500], goat anti-chicken IgY (H+L) [Abcam (ab150172), 1:500], rabbit anti-TGF-β (A15103), rabbit anti-Bcl-2 (A0208), and rabbit anti-Bax (A0207) were obtained from Abclonal (Beijing, China). Rabbit anti-cFOS [SYSY (226 004)], 1:200], donkey anti-chicken secondary antibody conjugated with Alexa Fluor 488, donkey anti-rabbit IgG (H+L) highly cross-adsorbed secondary antibody conjugated with Alexa Fluor 555, donkey anti-mouse secondary antibody conjugated with FITC 493 , and rabbit anti-goat secondary antibody conjugated with Cy3 were purchased from Abcam (USA). Donkey anti-rabbit secondary antibodies conjugated with HRP were purchased from Abclonal (Beijing, China). Adenovirus-associated virus, AAV2/9-mCherry, was generated by the viral core of Zhejiang University, and its titer was adjusted to 1 × 1013 copies per mL for injection.

### Characterization

Transmission electron microscopy (TEM) imaging was conducted using a Tecnai Spirit electron microscope at 120 KV (Thermo, Czech Republic). The hydrodynamic radius and zeta potential were characterized via dynamic light scattering (DLS) using a Litesizer 500 (Anton Paar, Austria). ^1^H-NMR spectra were obtained by an Avance III 500 M (Bruker, Germany). The averages of molecular weight (Mn) were characterized by gel permeation chromatography (GPC), which was performed on a modular system composed of a Waters 515 high-pressure liquid chromatographic pump, with DMF as the eluent (1 mL/min), and a Viscotek LR40 refractometer. All samples (5 mg) for analysis were dissolved in ∼1 mL of DMF and filtered through 0.2 μm filters prior to injection.

### Synthesis and characterization of GABA-CLP and DOPA-CLP

To synthesize GABA-CLP, CLP-257 (30.7 mg, 0.1 mmol, 1 equiv.) was dissolved in DMSO (2 mL), and then N,N’-carbonyl diimidazole (20 mg, ∼0.12 mmol, 1.2 eq.) was added to the solution. After 30 min, GABA (11 mg, 0.1 mmol, 1 eq.) was added to the mixture for the formation of modified CLP. The mixture was then precipitated several times in deionized water followed by washing several times with methanol for purification. Finally, the pale-yellow solids (GABA-CLPs) were collected after recrystallization or freeze drying and were characterized by ^1^H-NMR. To identify GABA-CLP’s targeting and metabolic process, fluorescent labeling was prepared. A similar procedure was used to prepare GABA-Cy5.5 using Cy5.5 instead of CLP-257.

DOPA-CLP was obtained by a similar synthetic procedure. Briefly, CLP-257 (30.7 mg, 0.1 mmol, 1 eq.) and N,N’-carbonyl diimidazole (20 mg, ∼0.12 mmol, 1.2 eq.) were dissolved in DMSO (2 mL). After 30 min, DOPA (19 mg, 0.1 mmol, 1 eq.) was added to the mixture, precipitated several times in deionized water, and then washed several times with toluene to obtain a pale yellow solid. After recrystallization, the product was named DOPA-CLP and was characterized by ^1^H-NMR. For fluorescent observation, DOPA-Cy5.5 was prepared by a similar procedure with Cy5.5 instead of CLP-257.

### Synthesis and characterization of copolymer (POEGMA-BAA)

The copolymers were prepared using reversible addition-fragmentation chain transfer (RAFT) polymerization. To optimize the molecular weight of the copolymer, four other kinds of micelles with copolymers of POEGMA_15_-BAA_2_, POEGMA_30_-BAA_2,_ POEGMA_40_-BAA_2_ and POEGMA_50_-BAA_2_ were prepared.

The representative synthetic procedure of POEGMA_30_-BAA_2_ was as follows: First, CTA (28 mg, 0.1 mmol, 1 eq.), OEGMA (1.66 g, 3.5 mmol, 35 equiv.)), AIBN (1.6 mg, 0.01 mmol, 0.1 equiv.) were dissolved in 2 mL DMF under a N_2_ atmosphere in a Schlenk tube. The mixture solution was degassed by three repeated freeze–evacuate–thaw cycles to remove the dissolved oxygen. The polymerization was carried out at 70°C for 12 h. The mixture was stopped by exposure to air and was precipitated in diethyl ether three times to obtain a red oil named POEGMA. Then, POEGMA (0.72 g, ∼1 eq.), 3-acrylamidophenylboronic acid (BAA, 3 mg, 0.016 mmol, 3 equ.), AIBN (0.8 mg, 0.005 mmol, 1 equ.) were dissolved in 1.5 mL DMF under a N_2_ atmosphere, frozen in liquid nitrogen and degassed for 30 min to remove dissolved oxygen. The polymerization was carried out at 70°C and was stopped after 10 h by exposure to air. The mixture was precipitated in diethyl ether three times to obtain copolymer POEGMA-BAA_2_ as an orange oil. The obtained POEGMA and POEGMA-BAA_2_ were characterized by ^1^H-NMR. Copolymers [POEGMA_15_-BAA_2_, POEGMA_30_-BAA_2,_ POEGMA_40_-BAA_2_ and POEGMA_50_-BAA_2_] with various molecular weights were obtained using different OEGMA and BAA monomer feeding ratios.

### Synthesis and characterization of micelles and nanoparticles

The micelles and nanoparticles were prepared by membrane dialysis43. The copolymer with different molecular weights (150 mg) was dissolved in 10 mL DMSO and pushed through a syringe pump to produce self-assembled micelles in 5 mL deionized water. Micelles were obtained by dialysis against water for 24 h using porous dialysis tubing (MWCO 3,500), which completely removed DMSO. As a result, micelles formed from POEGMA_30_-BAA_2_ with good dispersibility and appropriate size were used in the following experiments.

Then, we encapsulated the prodrug of GABA-CLP or DOPA-CLP in the micelles for sustained drug release. POEGMA_30_-BAA_2_ (100 mg), GABA-CLP or DOPA-CLP (50 mg) was dissolved in 10 mL DMSO. The mixture underwent the same procedure for the micelle formation and DMSO removal that we described above. The obtained nanoparticles with prodrugs were named GABA Nano or DOPA Nano, respectively. The GABA Nano and DOPA Nano that were used for fluorescence observation were prepared using GABA-Cy5.5 and DOPA-Cy5.5 for micelle formation instead of GABA-CLP and DOPA-CLP. In all, the following types of micelles were prepared: CLP-257-binding GABA-targeting micelles (GABA Nano), CLP-257-binding DOPA-targeting micelles (DOPA Nano), Cy5.5-binding GABA-targeting micelles (GABA Nano) and Cy5.5-binding DOPA-targeting micelles (DOPA Nano), and micelles formed by POEGMA_30_-BAA_2_ without drug loading were used as controls (ROS Nano). The critical micelle concentration (CMC), particle size, size distribution and zeta potential of the micelles and nanoparticles were determined by using DLS. The morphology of the micelles and nanoparticles was characterized by TEM.

### Release and degradation of ROS Nano by ROS stimulation

To characterize the features of ROS Nano under ROS stimulation, micelles (ROS Nano@Nile Red) were formed by POEGMA_30_-BAA_2_ (100 mg) and Nile red (40 mg) using the membrane dialysis method. Then, ROS Nano@Nile Red (∼5 mg, containing 1 mg Nile red) was dissolved in H_2_O_2_ solutions with different concentrations (0, 50, 100 and 500 mM, 1 mL. The fluorescence intensity was measured at certain intervals, and the release profiles of ROS Nano with ROS stimulation were obtained according to the changes in fluorescence intensity. The morphological changes in the ROS Nano in H_2_O_2_ solution (500 mM, 5 mg/mL) were characterized by a transmission electron microscope after oxidation for 1 h and 2 h.

### Biocompatibility characterization

PC12 cells were cultured in RPMI 1640 medium supplemented with 5% fetal bovine serum, 10% horse serum, 1% penicillin/streptomycin, and 2 mM l-glutamine for proliferation. A standard CCK-8 assay was used to test cell toxicity. Briefly, suspended PC12 cells were seeded in 96-well cell culture plates (10,000 cells per well) and cultured in medium for 24 h. Then, ROS Nano, GABA Nano or DOPA Nano was added to the medium at concentrations of 0, 31, 63, 125, 250, and 500 μg/mL, respectively, and the samples were maintained for 24 h. After washing twice with PBS, 100 μL of CCK-8 solution was added to each well and the samples were incubated for another 2 h. The absorbance of each well was measured with 450 nm laser excitation, and the cell survival ratio was obtained in relation to that in the control groups.

According to the manual, viability was further investigated with a Live/Dead viability/cytotoxicity kit. Briefly, cells treated with ROS Nano for 24 h were stained with 100 μL of 2 mM acetyl methoxy methyl ester (Calcein-AM) or 1.5 mM propidium iodide (PI) for 15 min at 37°C in 1× buffer and were then observed by inverted fluorescence microscopy (Olympus IX53, Japan).

### Cell protection from ROS damage by ROS-responsive nanodrugs

To assess whether ROS-responsive nanodrugs can protect cells from ROS-induced apoptosis, H2O2 (100 μM)-containing medium was used in cell culture. ROS Nano, GABA Nano or DOPA Nano was added to the medium at concentrations ranging from 0 to 1 mg/mL. After 24 h, the cells were washed twice with PBS and tested by CCK-8 assay.

Lipopolysaccharide (LPS)-containing medium was used to simulate the ROS microenvironment in damaged tissues. Cells were cultured with LPS (100 ng/mL)-containing medium and treated with ROS Nano, GABA Nano or DOPA Nano (250 μg/mL) for 6 h. Then, 2′,7′-dichlorofluorescin diacetate (DCFH-DA) detection was performed by adding 10 μM DCFH-DA for 20 min at 37°C followed by inverted fluorescence microscope observation.

### Surgical procedures

The spinal cord contusion injury model was established by an Infinite Vertical impactor (68099, RWD Life Science, China). To expose the dorsal surface of the spinal cord, laminectomy was performed at the 10th thoracic vertebral level (T10-11) after rats had been anesthetized with pentobarbital sodium (0.5 mL/100 g). Then, the impactor tip was lowered until it just contacted the exposed spinal cord. The spinal cord contusion was performed by smashing an impactor tip into the spinal cord 2.5 cm with a 3-mm diameter cylinder at a velocity of 2.5 m/s. Following surgery, the muscles and skin were sutured, and the rats were placed on an electric heating pad to maintain their body temperatures at 32°C until they awoke. Bladder care was provided twice daily until spontaneous voiding resumed. The rats used for the experiments were distributed into the following groups: the PBS group, ROS Nano group, GABA Nano group and DOPA Nano group. For the first week after SCI, 200 µL of each Nano solution (10 mg/mL) or PBS was administered by tail vein injection every 48 h. During the following weeks, 200 µL of each Nano solution (10 mg/mL) was administered via tail vein injection once a week.

A T10 complete crush model was performed according to the method described in previous literature6. Briefly, a midline incision was made over the thoracic vertebrae, followed by a T10 laminectomy. A complete T10 crush was then carefully conducted using surgical forceps (width, 0.1 mm). After surgery, the muscles and skin were sutured, and the rats were placed on an electric heating pad to maintain their body temperatures at 32°C until they awoke. All surgeries were completed by a surgical technician who was blinded to the experimental conditions.

To perform the anterograde tracing of propriospinal axons, rats underwent dorsal laminectomy at the 8^th^ thoracic vertebral level (T7-8 spinal cord). Then, according to the method described in previous literature50, AAV2/9-mCherry was injected into the T7-8 spinal cord of rats two weeks before termination. Histological assessment was performed 9 weeks after injury.

### Histology *and* 3D reconstruction

To collect spinal cord tissues, the rats were deeply anesthetized with pentobarbital sodium (0.5 mL/100 g) and perfused intracardially with 4% paraformaldehyde at the 9^th^ week postinjury. Before embedding, the fixed spinal cords were immersed in 30% and 15% sucrose solutions for dehydration. Spinal cord blocks were sectioned using a cryostat (CryoStar NX50; Thermo, USA) and thaw-mounted onto Super Frost Plus slides (Fisher Scientific, USA). Then, these sections were processed for immunohistochemistry assessments. After blocking with PBS with 5% donkey serum and 0.3% Triton X-100, spinal cord sections were incubated with primary antibodies. Then, the sections were washed three times with PBS and incubated with the appropriate secondary antibodies. Finally, the section slides were observed by confocal laser scanning microscopy (A1Ti, Nikon, Japan).

The quantitative analysis of the cavity volume of injured spinal cord tissues was performed according to a method described in previous literature50. Briefly, serial spinal cord cross sections equally spaced 200 μm apart were stained with hematoxylin and eosin (H&E) and imaged by a virtual digital slice scanning system (VS120, Olympus, Japan) for three-dimensional reconstruction. A 3D image was created by Amira software, and the cavity and pathological tissue were quantified.

### Toxicity and anti-inflammatory function of ROS Nano in vivo

To assess ROS nanotoxicity, rats treated with ROS Nano or PBS at the time points described previously were anesthetized and perfused with 4% paraformaldehyde on the 7^th^ day postinjury. Sections of the heart, liver, spleen, lung, kidney and brain were collected and subjected to H&E staining for the observation of tissue morphologies. To assess the anti-inflammatory function of ROS Nano, spinal cord segments 1-2 mm rostral and caudal to the injury site were dissected. Then, the tissue was homogenized using a homogenate in lysis buffer and centrifuged at 12,000 g and 4°C for 10 min to obtain the supernatant. The total protein content was measured with a BCA protein assay kit. Forty micrograms/sample protein was loaded onto a 10% polyacrylamide gel, separated by SDS PAGE, and transferred to polyvinylidene fluoride membranes. After blocking with 5% skimmed milk in PBST for 1 h at room temperature, membranes were placed in primary antibody (rabbit anti-TGF-β, rabbit anti Bcl-2, rabbit anti Bax, 1:1000 dilution by 5% BSA) overnight at 4°C. After washing with PBST 3 times, the membrane was incubated with a secondary antibody (goat anti-rabbit HRP) for 1 h at room temperature. Then, protein signals were visualized using an ECL kit and measured by Image Lab Software provided by Bio–Rad.

### Drug targeting and distribution in vivo

To assess the targeting effect and biodistribution of nanodrugs, GABA Nano and DOPA Nano were administered via tail vein injection at 3 h postinjury. At certain intervals (after injection for 3, 6, 24 and 48 h), the rats were deeply anesthetized with pentobarbital sodium (0.5 mL/100 g) and perfused intracardially with 4% paraformaldehyde. The heart, liver, spleen, lung, kidney, brain and spinal cord were collected and observed with an in vivo fluorescence imaging system (CRi Corporation, America, MK50101-EX). To assess the BSCB penetration capacity of nanodrugs, ROS nanoinjection was performed in SCI rats at 4, 7, and 14 days postinjury or intact rats. Spinal cords were dissected at 6 h postinjection and assessed by both an in vivo fluorescence imaging system and immunofluorescence histology.

### Behavioral assessment

Behavioral assessment of rats was performed weekly in an open-field environment based on the original report of the BBB51. Rats that showed a BBB score above 1.5 at 1 week following contusion were excluded from further studies. To analyze the detailed hindlimb kinematics, procedures were performed according to the method described in previous literature50. The hindlimb movement of rats from different groups was recorded using the MotoRater (Vicon Motion Systems, UK)52. The stick views of the hindlimb and angle of rotation movement were obtained by MATLAB in a double-blinded manner.

The behavioral data were depicted with seven features (maximal iliac crest height, crest height amplitude, maximal toe height, toe height amplitude, hip, knee and ankle angle oscillation) within five groups (intact, PBS, ROS Nano, DOPA Nano, and GABA Nano). Because we utilized different statistical methods to calculate the seven behavioral features, these features also had different units. To depict these features together in the radar graph, we normalized the seven behavioral feature data with equation (1):

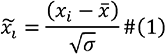

where *x_i_* serves as the *ith* data in each of the seven behavioral features, *i* = 1,2,3,…*n*. . *n* is the number of points sampled from each feature in each group. *x̄* and √σ serve as the mean value and standard variance in each feature, respectively. In the next step, we obtained seven *x̃_i_* values with both positive and negative values in each group. We then projected the features into an exponentiation expression and mapped these values in a unified scale (a unified vector space). For each depicted feature, we use equation (2) to process the data:

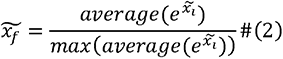

where *f* = 1,2,3,…,7, serves as the seven behavioral features in each group. We averaged the 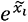 in each group and obtained a list with seven feature values in each group.

Finally, we used the intact group values as the benchmark and depicted the feature values’ variance. We used equation (3) to process the data:

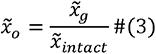

where *x̃_g_* serves as each feature value in each group. *x̃_intact_* serves as the feature values in the intact group, namely, the benchmark values.

### Electromyography (EMG) recording and data analysis

At 9 weeks after SCI, the implantation of bipolar electrodes was performed according to a method described in previous literature50. Briefly, electrodes (AS632, Conner wire) were led by 25-gauge needles and inserted into the mid-belly of the medial gastrocnemius (GS) and tibialis anterior (TA) muscles of one hindlimb when rats were deeply anesthetized. A common ground wire was inserted subcutaneously into the Achilles tendon area of the hindlimb. Wires were routed subcutaneously through the back to a small percutaneous connector securely cemented to the skull of the rat. EMG signals were acquired using a differential neuron signal amplifier (BTAM01 L, Braintech, China) with 30- to 2000-Hz filtration, sampled at 30 kHz using a Neurostudio system (Braintech, China), and analyzed by a custom MATLAB code.

In this study, the Poincaré analysis method was used to distinguish chaos from randomness by embedding data sets into a higher dimensional state space. For example, a time series of EMG peak signals was given as follows:

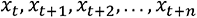

To obtain a return map in its simplest form, we plotted (x_t_, x_(t+1)_), (x_(t+1)_, x_(t+2)_), and then (x_(t+2)_, x_(t+3)_), and so forth. In this way, the interval length between two EMG peak signals in a series could be clearly represented in the plot, facilitating rhythm quantification among different groups (Intact, PBS, ROS Nano, GABA Nano, and DOPA Nano). Finally, we obtained both amplitude and rhythm information by analyzing the EMG data.

### Statistical analysis

Multiple samples were analyzed by one-way or two-way ANOVA. Differences between the vehicle control and experimental groups were analyzed by Student’s t test. A P value less than 0.05 was considered significant.

## Acknowledgments

The authors acknowledge the excellent technical staff at the Imaging Facility, Core Facility of Zhejiang University School of Medicine and Center of Cryo-Electron Microscopy of Zhejiang University for their assistance with confocal laser scanning microscopy and scanning electron microscopy. This study was supported by the National Natural Science Foundation of China (81971866 to X.W.), the Natural Science Foundation of Zhejiang Province (LR20H090002 to X.W.), Fundamental Research Funds for the Central Universities (K20210195 to X.W.), and the National Major Project of Research and Development (2017YFA0104701 to B.Y.).

## Author Contributions

Y.Z. and J.Y. contributed equally to this work. X.W. and Y.Z. conceptualized and designed the study. Y.Z., J.Y., W.C., X.C., L.L., B.G., S.J., H.Z., A.F., X.Q. and X.W. conducted the experiments. Y.Z. J.Y. and W.C. collected the data. Y.Z., J.Y., W.C., B.G., X.G., Z.W., Z.Z., B.Y., and X.W. analyzed and interpreted the data. Y.Z., X.W., J.Y. and Z.A. drafted the paper. All authors critically revised the manuscript and approved the final version for submission.

## Data Availability Statement

The data to support the findings of this study are included in the paper, and further data are available from the corresponding author upon reasonable request.

**Movie S1.** Vicon system observation of the movement of the rat hindlimb under 5 conditions (intact, PBS, ROS Nano, DOPA Nano, GABA Nano).

---

**Fig. S1.**
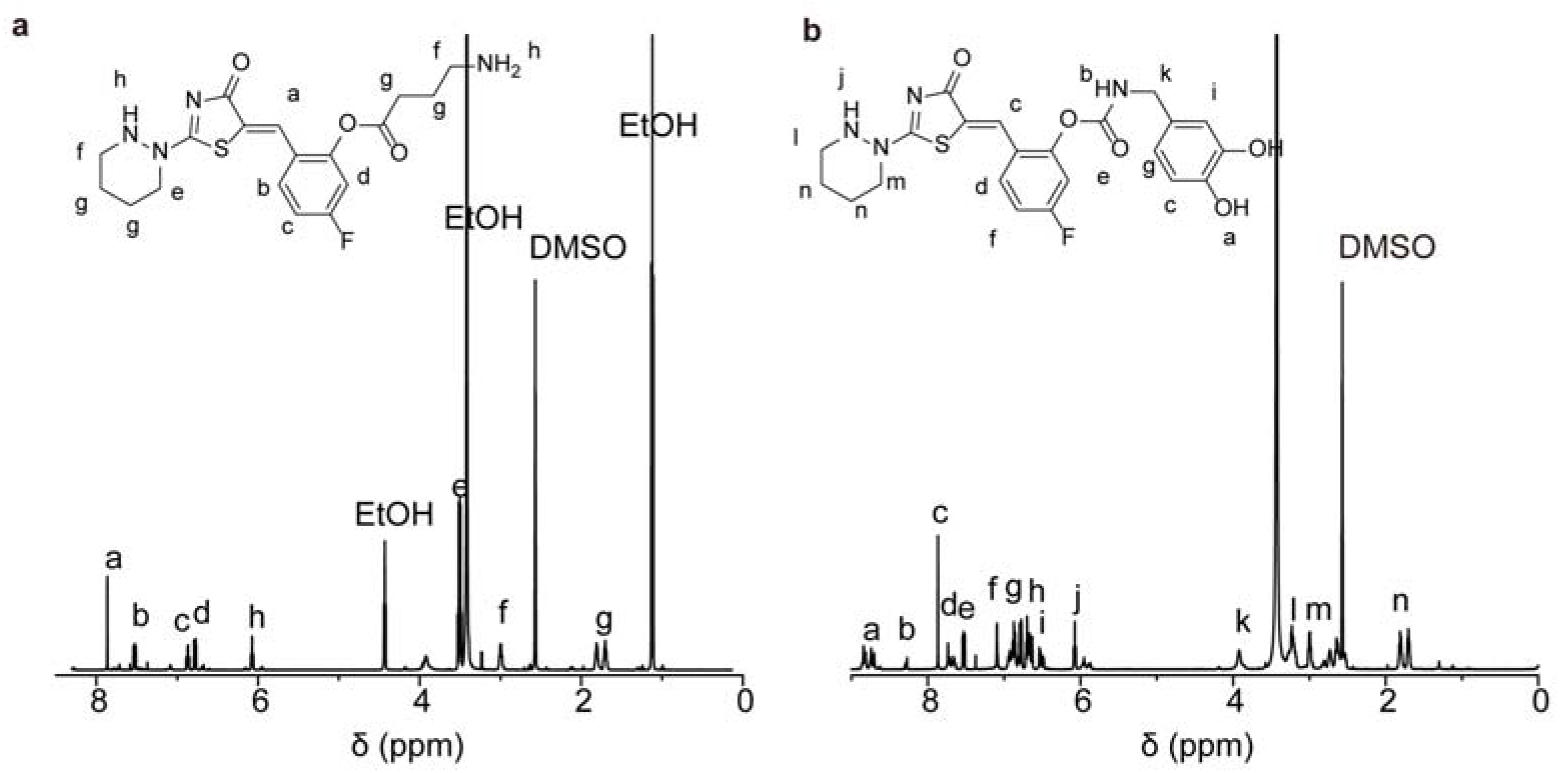
Characterization of different neurotransmitter functionalized modulators. **a-b**, the ^1^H-NMR spectra of GABA and dopamine-functionalized CLP-257.

**Fig. S2.**
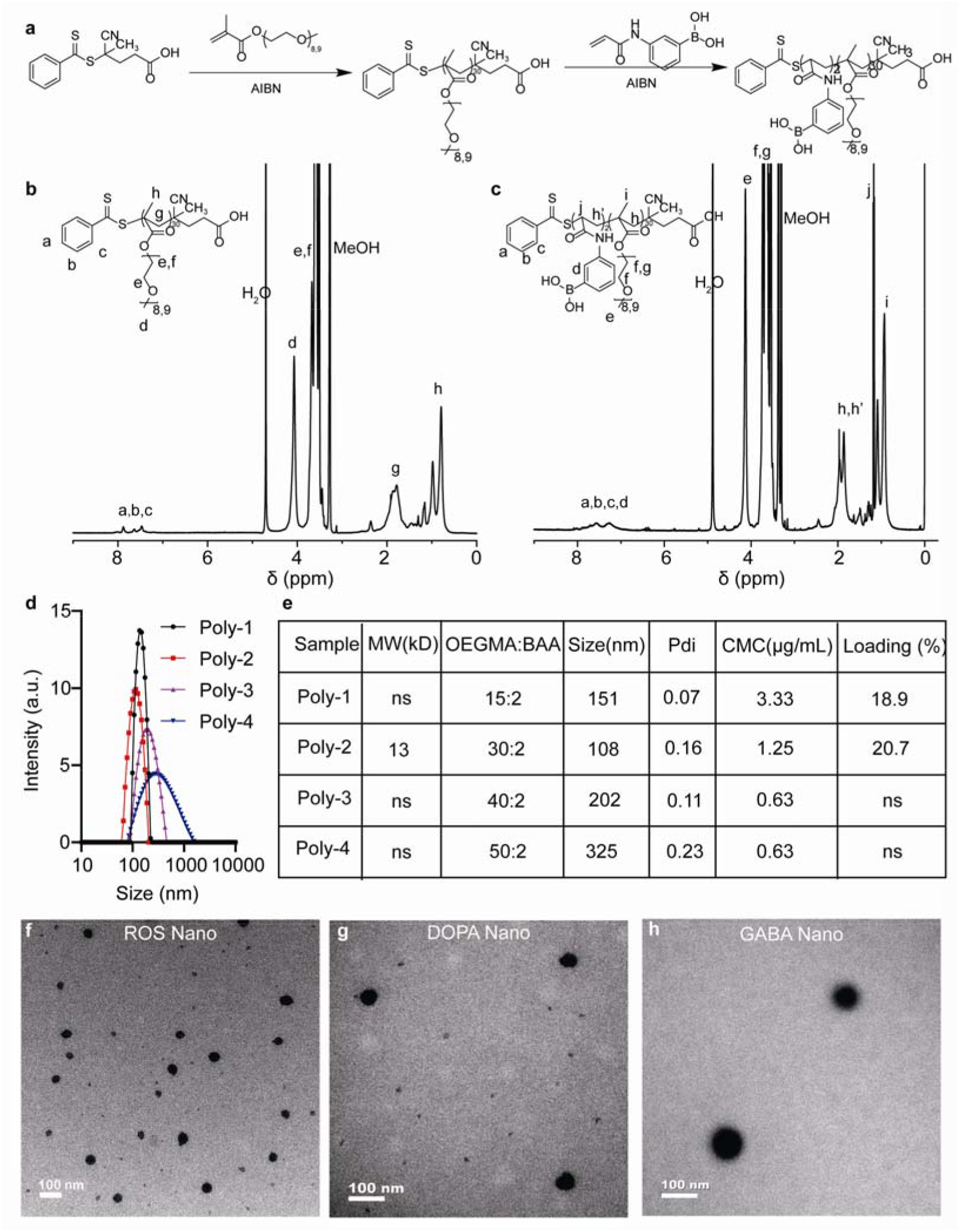
Synthesis and characterization of amphiphilic polymers and nanoparticles. **a**, The synthetic route of a representative amphiphilic polymer. **b-c**, Representative ^1^H-NMR spectra of POEGMA_30_ and POEGMA_30_-BAA_2_. **d-e**, Properties of series of nanoparticles obtained from different composition of amphiphilic polymers. **f-h**, Representative TEM images of ROS Nano, DOPA Nano and GABA Nano. Scale bar, 100 nm.

**Fig. S3.**
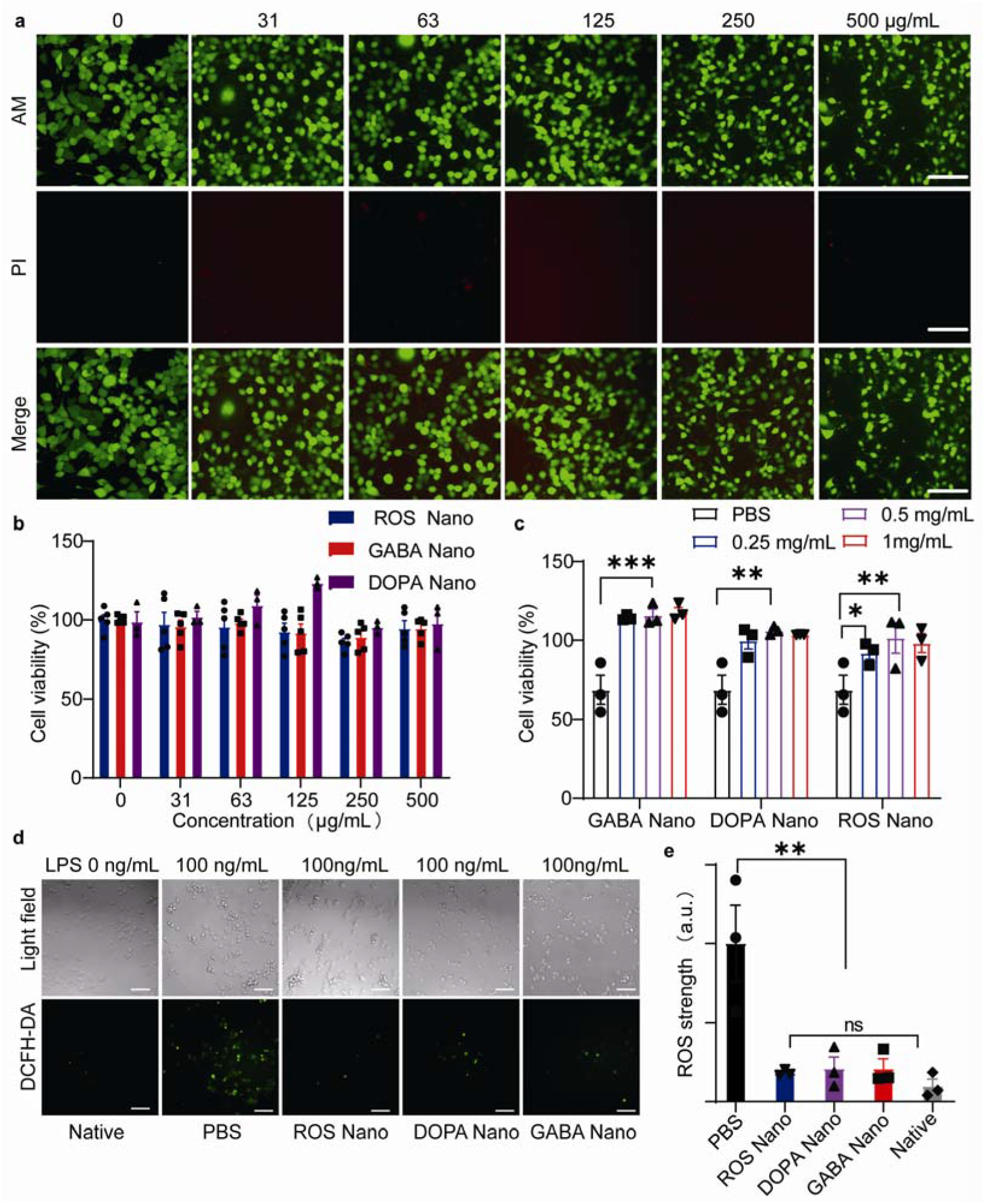
Cell compatibility and antioxidant properties of nanodrugs. **a**, Representative fluorescent images of PC12 cells after incubation with different concentrations of ROS Nano for 24 h. Calcein-AM (acetyl methoxy methyl ester) and PI (propidium iodide). Scale bar, 100 μm. **b**, Cell viability after incubation with different concentrations of ROS Nano, GABA Nano and DOPA Nano for 24 h. Data are shown as mean ± SEM. One-way ANOVA with Tukey’s post hoc test for comparisons among multiple groups. n = 3. *P < 0.05, **P < 0.01, ***P < 0.001 statistical significance. **c,** Cell viability after treatment with H_2_O_2_ solution (100 μM) for 24 h after incubation with different concentrations of ROS Nano, GABA Nano and DOPA Nano. Data are shown as mean ± SEM. One-way ANOVA with Tukey’s post hoc test was used for comparisons among multiple groups. n = 3 to 5. *P < 0.05, **P < 0.01, ***P < 0.001 statistical significance. **d**, Cellular ROS production of PC12 cells receiving different treatments after stimulation with LPS (100 ng/mL) determined by using DCFH-DA (2′,7′-dichlorofluorescin diacetate). Scale bar, 100 μm. **e**, Quantitative analysis shown in the bar graph. Data are shown as mean ± SEM. One-way ANOVA with Tukey’s post hoc test for comparisons among multiple groups. n = 3. *P < 0.05, **P < 0.01, ***P < 0.001 statistical significance.

**Fig. S4.**
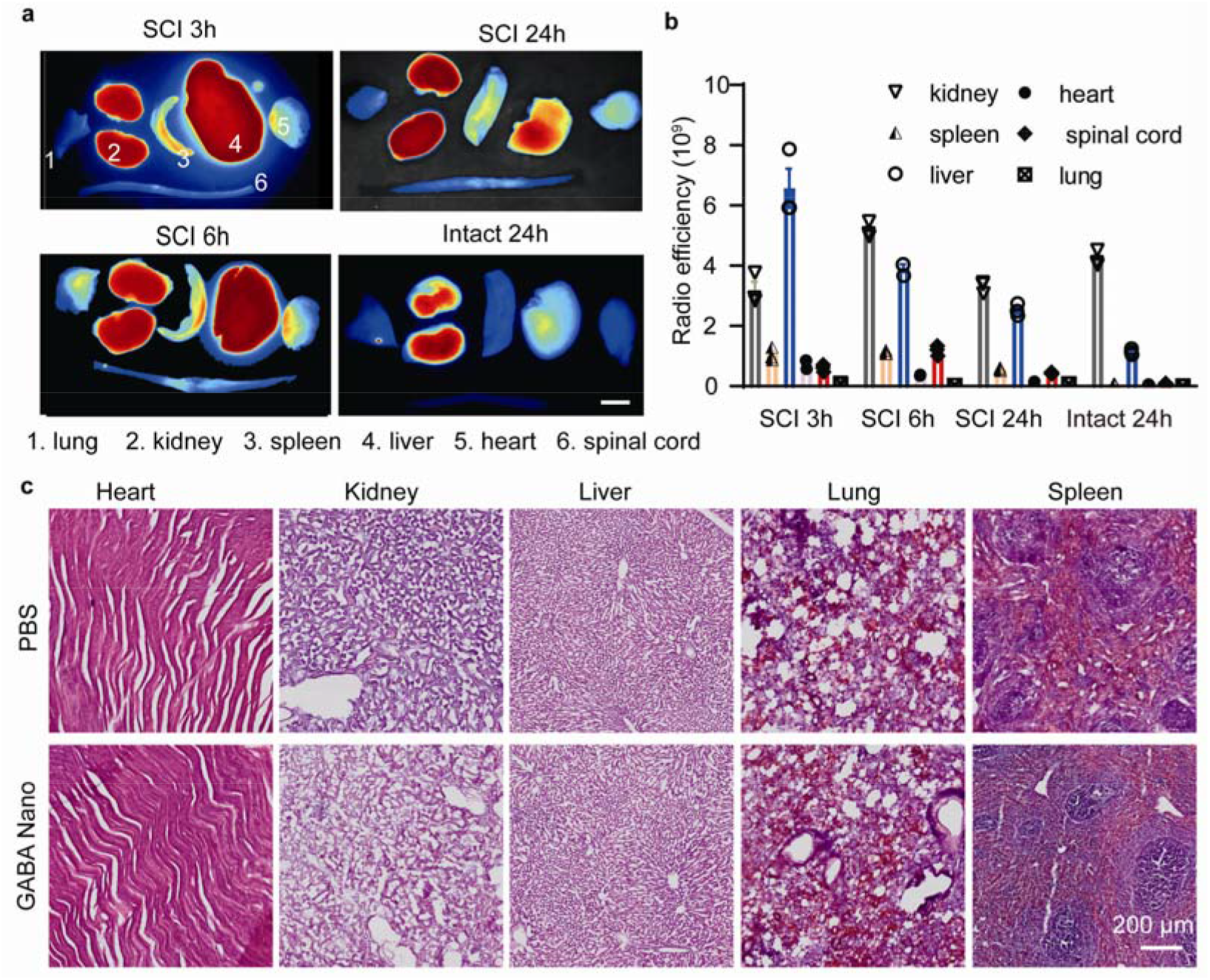
In vivo biodistributions of the nanodrugs after intravenous administration. **a**, Representative images of the main organs of rats that received a tail vein injection of ROS Nano@Cy5.5. Scale bar, 1 cm. **b**, Quantitative analysis of biodistributions in main organs of rats that received a tail vein injection of ROS Nano@Cy5.5. n = 3. Data are shown as mean ± SEM. **c**, Representative images of H&E-stained main organs sections after one week with ROS Nano (10 mg/kg) injection. Scale bar, 200 μm.

**Fig. S5.**
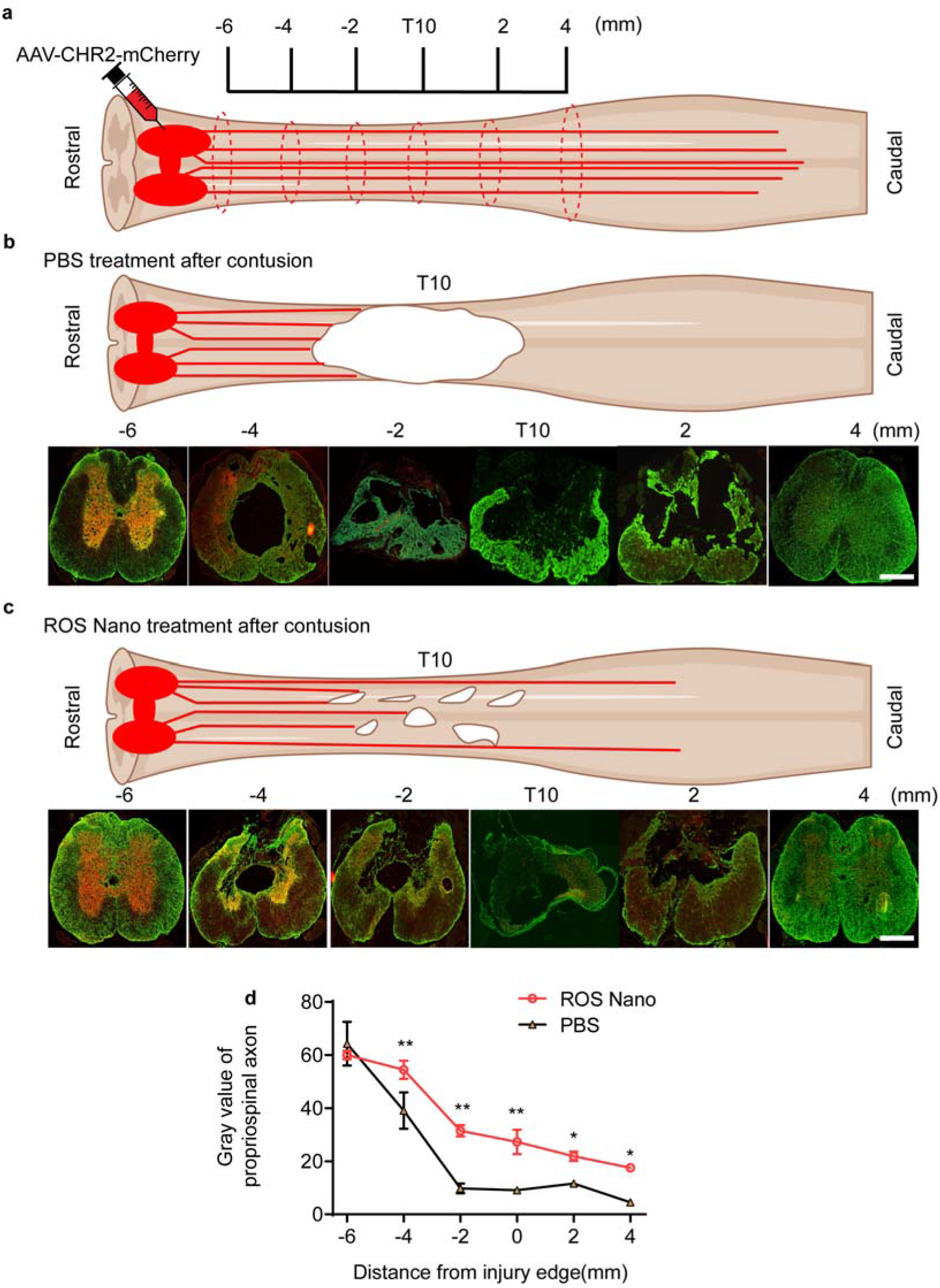
ROS nano treatment protects spared propriospinal axons from secondary injury. **a**, Schematic diagram of the experimental design. **b-c,** Representative images of cross sections at certain sites stained by RFP (representing propriospinal axon, red) and GFAP (green) in PBS or ROS Nano-treated rats. **d**, Quantitative analysis of propriospinal axons of rats with 9-week ROS Nano or PBS treatment. n = 3. Data are shown as mean ± SEM. One-Way ANOVA with Tukey’s post hoc test was used for comparisons among multiple groups. * and ** indicate p < 0.05 and p < 0.01, respectively.

**Fig. S6.**
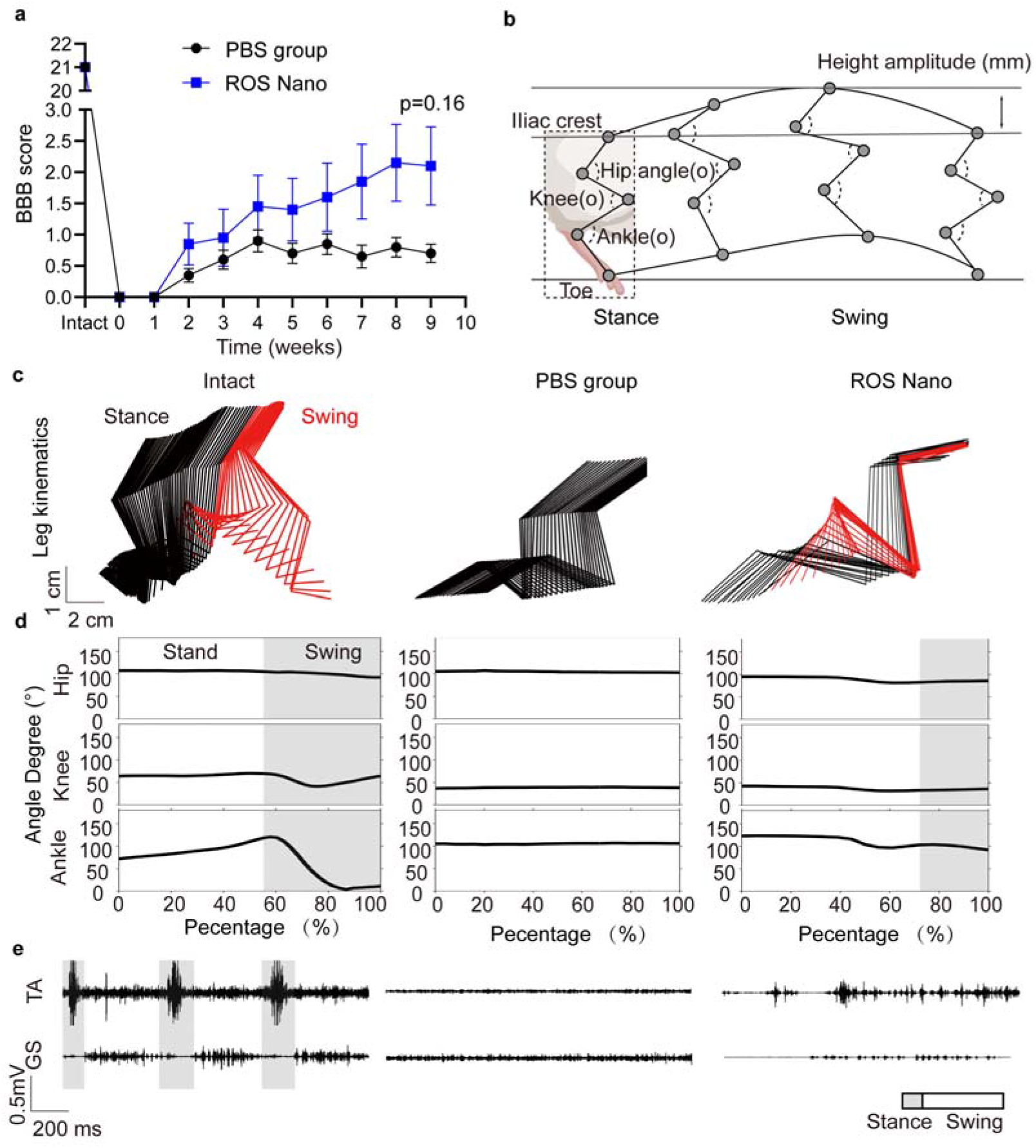
The rescued spared spinal connections mediate limited hindlimb locomotor functional recovery. **a,** Weekly BBB scores of the experimental rats treated with the PBS or ROS Nano treatment.; Data are shown as mean ± SEM. One-Way ANOVA with Tukey’s post hoc was used test for comparisons among multiple groups, two-tailed paired t tests were used for comparisons between two groups. n = 10. * and ** indicate p < 0.05 and p < 0.01, respectively. **b**, A simplified stick mode is illustrated to show a one-step cycle of one intact hindlimb while the rat walked freely. The model also shows the quantification of hip, knee and ankle angles and iliac crest height amplitudes during the cycle. **c**, Representative color-coded stick views of kinematic hindlimb movement of intact, PBS-treated or ROS Nano-treated rats. **d**, Representative curves of hip, knee and ankle angles during the one-step cycle. **e,** Representative EMG of the TA and GS muscles of rats with different treatments. Gray bars, stance; white bars, swing.

**Fig. S7.**
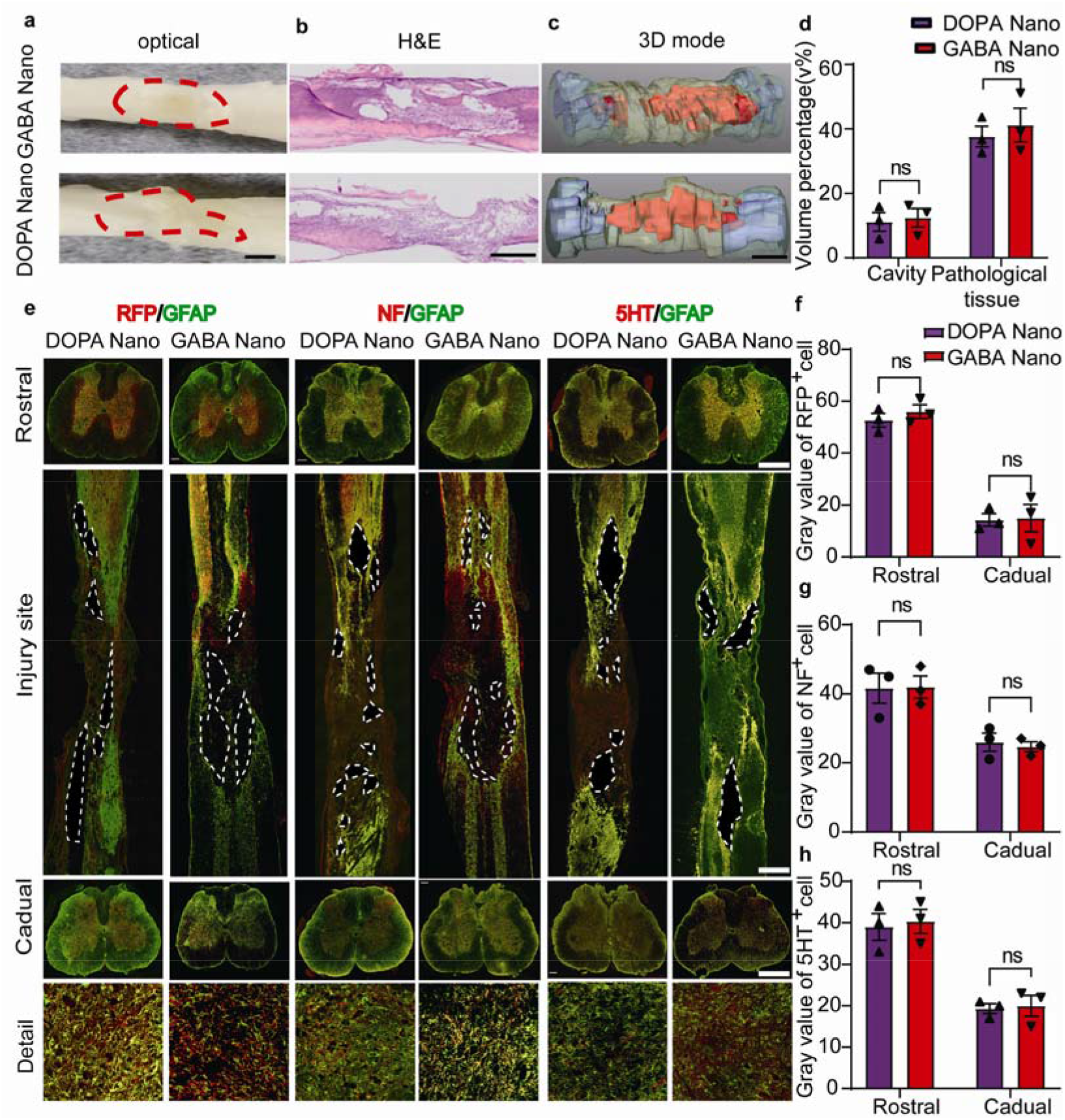
Spared tissues/axons in the spinal cord of rats with different treatments. **a**, Representative optical images of the spinal cords of rats treated with 9 weeks of GABA Nano or DOPA Nano treatment. A red dotted line indicates the boundary of injury. Scale bar, 1 cm. **b**, Representative H&E-stained images of the longitudinal sections of injured spinal cords. Scale bar, 1 mm. **c**, 3D reconstruction images of the spinal cord tissues with white matter (green), gray matter (light blue), pathological tissue (yellow), and cavities (red). Scale bar, 1 mm. **d**, Graph showing the quantification of the cavity and pathological volume percentages in injury sites. **e**, Representative images of the spinal sections stained with GFAP (green), NF (red), 5-HT (red) and RFP (red) in rats treated with GABA Nano or DOPA Nano for 9 weeks. Scale bar, 100 μm or 500 μm (middle). A white dotted line indicates the boundary of the cavity. rostral (upper), longitudinal (middle) and caudal (lower) spinal cord sections. **f-h**, Quantification of the RFP, NF and 5-HT immunoreactivity gray values on the rostral and caudal sides. Data are shown as mean ± SEM. One-way ANOVA with Tukey’s post hoc test was used for comparisons among multiple groups, and two-tailed paired t tests were used for comparisons between two groups. n = 3.

**Fig. S8.**
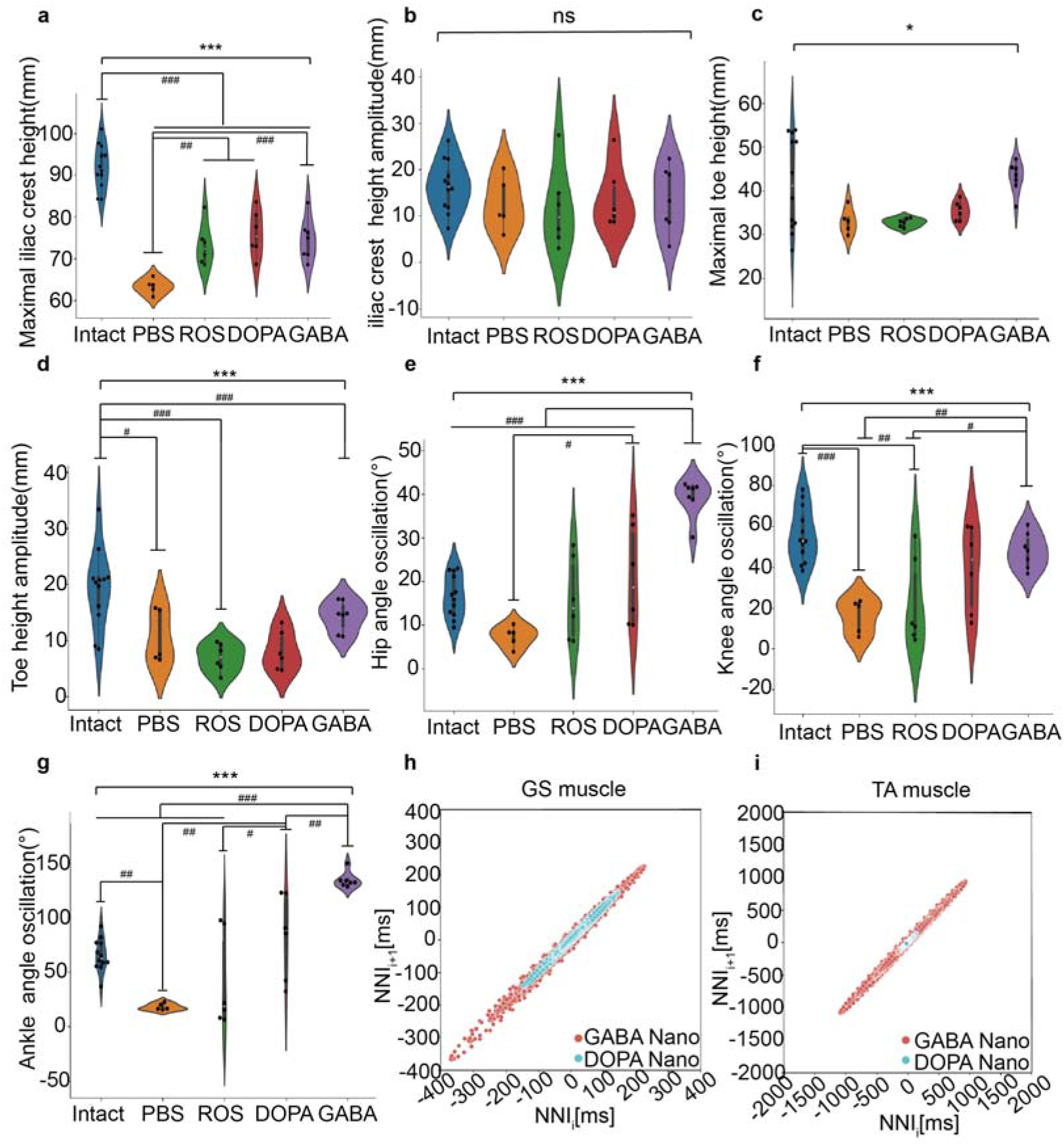
Detailed statistical analysis of the hindlimb movement of rats with different treatments. **a,** Quantification of the average maximal iliac crest height among groups. **b**, Quantification of the iliac crest height amplitude among the rats with different treatments. One-way ANOVA with Tukey’s post hoc test for comparisons among multiple groups (*) and two-tailed paired t tests were used for comparisons within groups (#) for the data shown in the violin plot. n = 5-12. ***p (or ###p) < 0.001, **p (or ##p) < 0.01, *p (or #p) <0.05. **c**, Quantification of the average maximal toe height among groups. **d,** Quantification of the average toe height amplitude among the rats with different treatments. One-way ANOVA with Tukey’s post hoc test for comparisons among multiple groups (*)and two-tailed paired t tests were used for comparisons within groups (#) for the data in the violin plot. n = 5-12. ***p (or ###p) < 0.001, **p (or ##p) < 0.01, *p (or #p) < 0.05. **e,** Quantification of the average hip oscillation among the rats with different treatments. One-way ANOVA with Tukey’s post hoc test for comparisons among multiple groups (*) and two-tailed paired t tests were used for comparisons within groups (#) for the data in the violin plot. n = 5-12. ***p (or ###p) < 0.001, **p (or ##p) < 0.01, *p (or #p) < 0.05. **f,** Quantification of the average knee angle oscillation among the rats with different treatments. One-way ANOVA with Tukey’s post hoc test for comparisons among multiple groups (*) and two-tailed paired t tests were used for comparisons within groups (#) for the data shown in the violin plot. n = 5-12. ***p (or ###p) < 0.001, **p (or ##p) < 0.01, *p (or #p) < 0.05. **g,** Quantification of the average ankle angle oscillation among the rats with different treatments. One-way ANOVA with Tukey’s post hoc test for comparisons among multiple groups (*) and two-tailed paired t tests were used for comparisons within groups (#) for the data in the violin plot. n = 5-12. ***p (or ###p) < 0.001, **p (or ##p) < 0.01, *p (or #p) < 0.05. **h-i,** Poincaré statistical analysis of the EMG signal amplitude rhythm of TA and GS muscles from DOPA Nano and GABA Nano groups. One-way ANOVA with Tukey’s post hoc test for comparisons among multiple groups (*) and two-tailed paired t tests were used for comparisons within groups (#) for the data shown in the violin plot. n = 5-12. ***p (or ###p) < 0.001, **p (or ##p) < 0.01, *p (or #p) < 0.05.

**Fig. S9.**
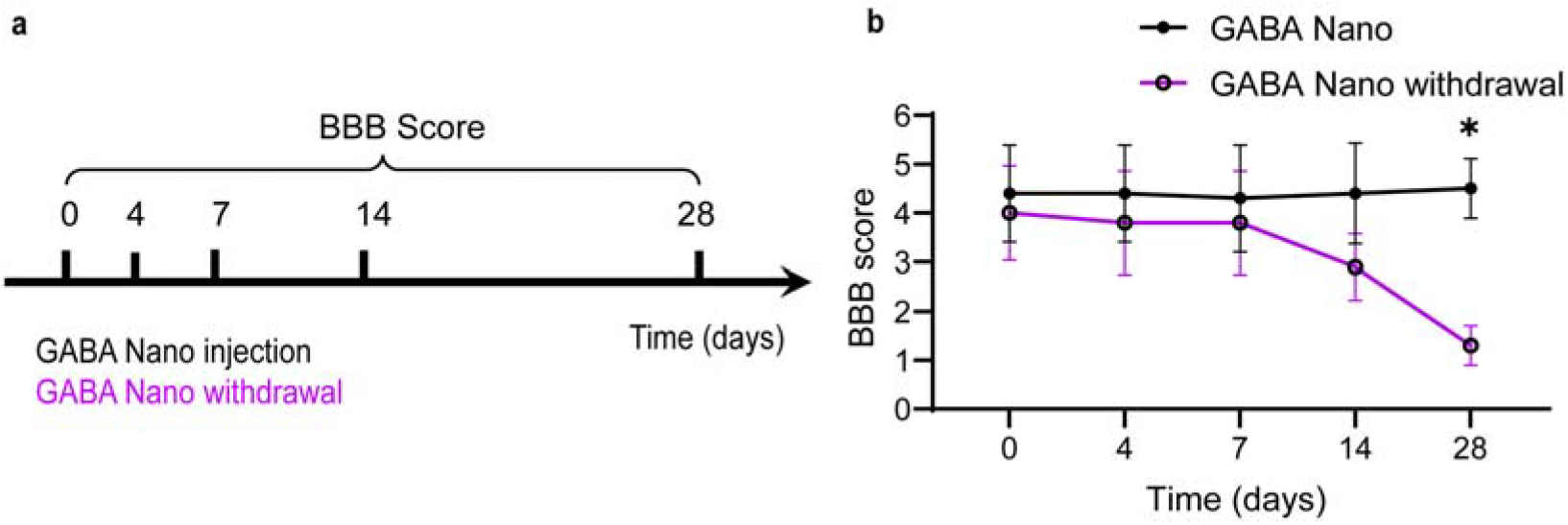
Behavioral changes after GABA Nano withdrawal in rats after 9 weeks of treatment. **a**, Schematic diagram of the experimental design of the GABA Nano and GABA Nano withdrawal groups. **b**, Weekly BBB scores of rats treated with PBS or GABA Nano. Data are shown as mean ± SEM. Two-tailed paired t tests were used for comparisons of two groups. n = 5. * Indicates p < 0.05.

**Fig. S10.**
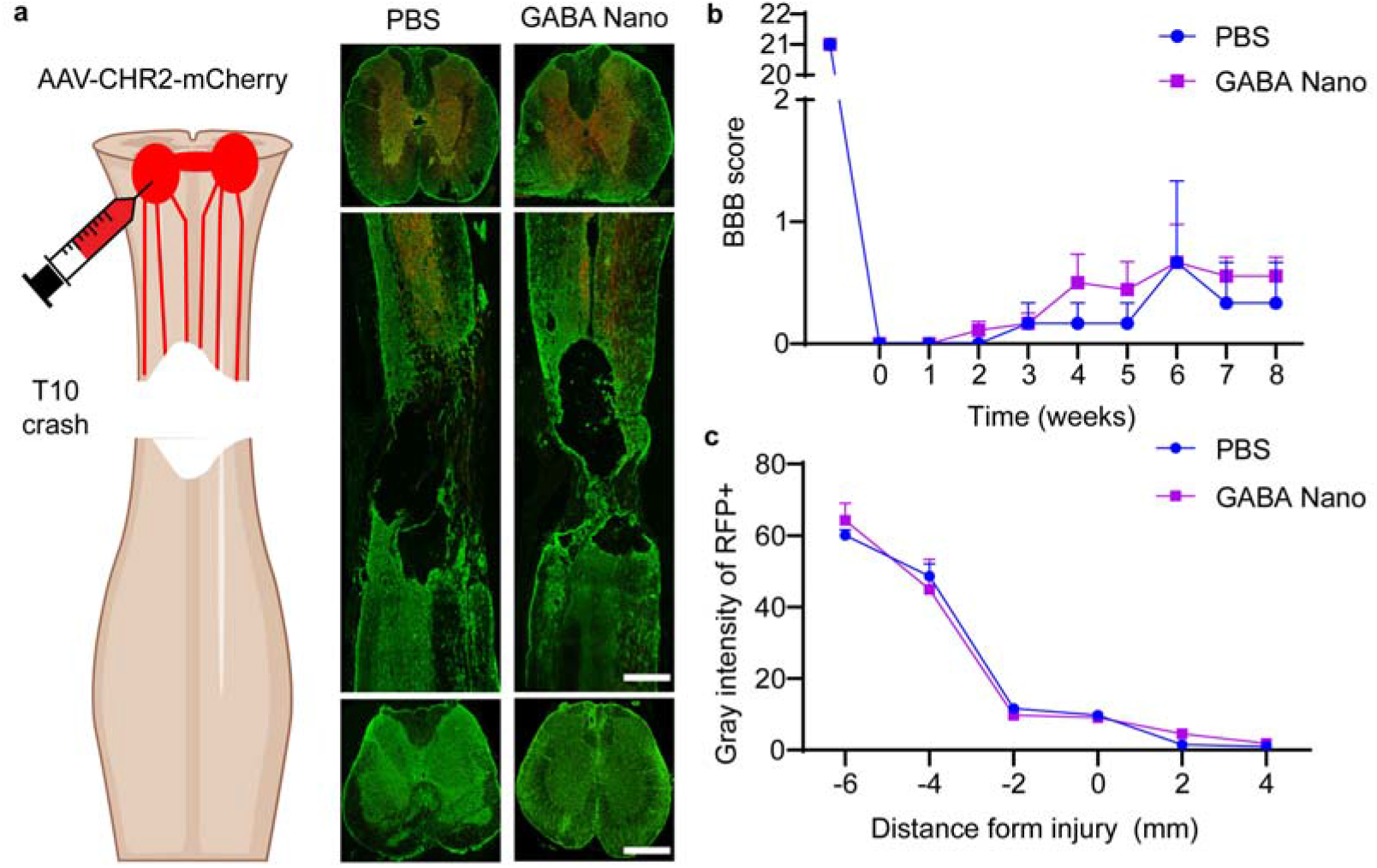
GABA Nano treatment shows no effect in rats with complete spinal cord Injury. **a**, Schematic diagram of the experimental design and representative images of longitudinal sections stained with RFP (red, representing propriospinal axons) and GFAP (green) of rats treated with PBS or GABA nano (10 mg/kg). **b**, Weekly BBB scores of rats treated with PBS or GABA Nano. Data are shown as the means + SEM. One-way ANOVA with Tukey’s post hoc test was used for comparisons among multiple groups, and two-tailed paired t tests were used for comparisons between two groups. n = 9. **c**, Quantitative analysis of the propriospinal axons in the spinal cord of rats with 8-week GABA Nano or PBS treatment. Data are the means + SEM. One-way ANOVA with Tukey’s post hoc test was used for comparisons among multiple groups, and two-tailed paired t tests were used for comparisons between two groups. n = 3. Scale bar, 100 μm.

**Fig. S11.**
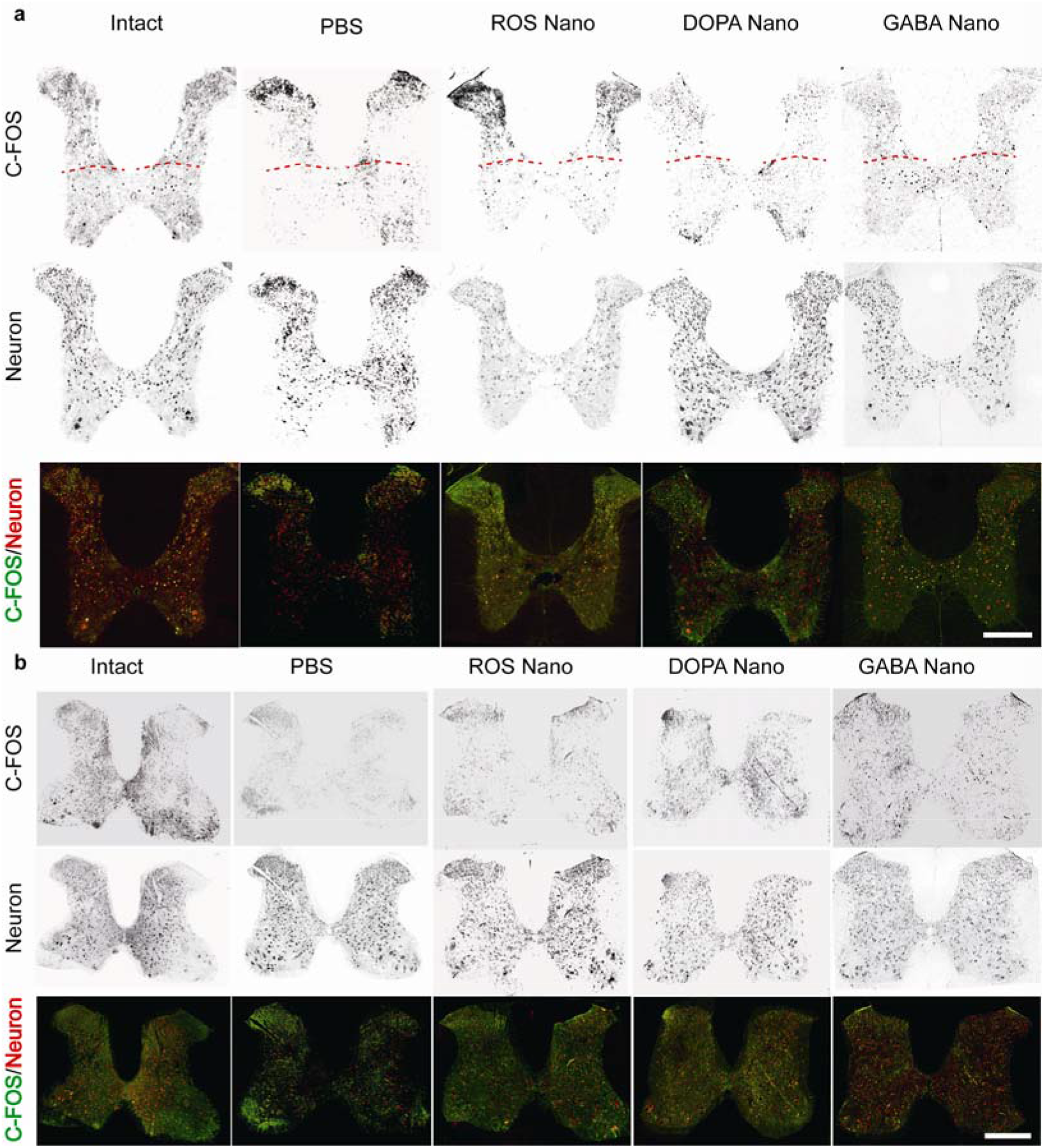
C-Fos expression in sections of T8 and L2 spinal cord with different treatments. **a**, Representative images of transverse sections stained with c-Fos and NeuN from the T8 spinal cord of injured rats with 9-week treatments of PBS, ROS Nano, DOPA Nano or GABA Nano. Scale bar, 100 μm. **b**, Representative images of the transverse sections stained with c-Fos and NeuN from the L2 spinal cord of injured rats after 9 weeks of treatment with PBS, ROS Nano, DOPA Nano or GABA Nano. Scale bar, 100 μm. T8 represents the 7^th^ thoracic vertebral level, and L2 represents the second Lumbala’s vertebral level.

